# Demography, education, and research trends in the interdisciplinary field of disease ecology

**DOI:** 10.1101/2020.07.16.207100

**Authors:** Ellen E. Brandell, Daniel J. Becker, Laura Sampson, Kristian M. Forbes

## Abstract

Micro- and macroparasites are a leading cause of mortality for humans, animals, and plants, and there is great need to understand their origins, transmission dynamics, and impacts. Disease ecology formed as an interdisciplinary field in the 1970s to fill this need and has recently rapidly grown in size and influence. Because interdisciplinary fields integrate diverse scientific expertise and training experiences, understanding their composition and research priorities is often difficult. Here, for the first time, we quantify the composition and educational experiences of a subset of disease ecology practitioners and identify topical trends in published research. We combined a large survey of self-declared disease ecologists with a literature synthesis involving machine-learning topic detection of over 18,500 disease ecology research articles. The number of graduate degrees earned by disease ecology practitioners has grown dramatically since the early 2000s. Similar to other science fields, we show that practitioners in disease ecology have diversified in the last decade in terms of gender identity and institution, with weaker diversification in race and ethnicity. Topic detection analysis revealed how the frequency of publications on certain topics have declined (e.g., HIV, serology), increased (e.g., the dilution effect, infectious disease in bats), remained relatively common (e.g., malaria ecology, influenza, vaccine research and development), or have consistently remained relatively infrequent (e.g., theoretical models, field experiments). Other topics, such as climate change, superspreading, emerging infectious diseases, and network analyses, have recently come to prominence. This study helps identify the major themes of disease ecology and demonstrates how publication frequency corresponds to emergent health and environmental threats. More broadly, our approach provides a framework to examine the composition and publication trends of other major research fields that cross traditional disciplinary boundaries.

## INTRODUCTION

Infectious diseases are a leading source of human, domestic animal, and wildlife mortality, killing an estimated 17 million people each year (Brand, 2013; World Health Organization 1996, 2018), threatening economic security through crop and production animal losses (Ahmed et al., 2019; Benavides, Rojas Paniagua, Hampson, Valderrama, & Streicker, 2017), and causing declines of endangered species (Scheele et al., 2019). Moreover, infectious disease outbreaks are predicted to be exacerbated by contemporary issues such as climate change, high human population density, and fragmentation of natural environments (Altizer, Ostfeld, Johnson, Kutz, & Harvell, 2013; Daszak, Cunningham, & Hyatt, 2001; Plowright et al., 2021). It is important to establish a strong foundation and specialization for research on infectious diseases in their ecological and evolutionary context to promote a high standard of living and enhance wildlife and ecosystem health; for example, we continue to face challenges such as emerging pathogens (e.g., SARS-CoV-2; Andersen, Rambaut, Lipkin, Holmes, & Garry, 2020), pathogen evolution (van Boeckel et al., 2019), and the need for innovative interventions (Sokolow et al., 2019).

Disease ecology is the study of how micro- and macroparasites move through and are distributed across host populations, landscapes, and ecosystems, considering both abiotic and biotic factors, as well as the consequences of their infections. It is a relatively new and rapidly expanding research focus within ecology and evolutionary biology that draws heavily on early foundations in population biology (Anderson & May, 1979; May & Anderson, 1979) and vector-borne disease (e.g., zooprophylaxis as a precursor to the dilution effect literature; (Hess & Hayes, 1970; Schmidt & Ostfeld, 2001). Further, disease ecology integrates many fields that cross multiple levels of biological organization including but not limited to parasitology, microbiology, immunology, and epidemiology (Grenfell, Dobson, & Moffatt, 1995; Hudson, Rizzoli, Grenfell, Heesterbeek, & Dobson, 2002; Wilson, Fenton, & Tompkins, 2019). Disease ecologists investigate a range of practical and fundamental questions relevant to humans, other animals, and plants, such as the natural origins of disease outbreaks; heterogeneities in pathogen susceptibility, transmission, and impact; and the effectiveness of intervention strategies (Condeso & Meentemeyer, 2007; Hudson, Dobson, & Newborn, 1998; Joseph et al., 2013; Olival et al., 2017; Vanderwaal & Ezenwa, 2016).

Disease ecology, in part, adapted and developed population biology theory to address societal needs (Johnson, de Roode, & Fenton, 2015; Koprivnikar & Johnson, 2016; Scheiner & Rosenthal, 2006). Key among these is the urgency to understand and address novel disease threats, which are rooted in natural systems but are often exacerbated by societal inequalities (Carlson & Mendenhall, 2019). For example, the impacts of habitat degradation on pathogen spillover is an expanding area of research that can be used to guide risk assessments and environmental policy (Plowright et al., 2021). At the same time, infrastructure has developed around disease ecology, including journals and associated organizations (e.g., Wildlife Disease Association, American Society of Tropical Medicine and Hygiene), and a specialized National Science Foundation and National Institutes of Health funding program and conference series (Scheiner & Rosenthal, 2006), which have helped to direct research effort and create networks amongst researchers.

Still, many questions remain as to the composition of disease ecology practitioners, core research foci, and if research trends are associated with widespread disease outbreaks. Answering these questions could help improve recruitment and retention and prioritize future research directions. However, understanding these complex and interrelated factors as they apply to an interdisciplinary research field requires diverse and innovative approaches. Here we characterize the field of disease ecology and a subset of its practitioners by addressing the following questions: (1) Who comprises the field in terms of education, demographics, and the type of research they conduct? (2) Which scientific articles and journals have been the most influential? (3) And significantly, how has the frequency of research topics emerged and changed in the literature over time? For example, do the topics in publications follow global health events such as disease outbreaks?

To answer these questions, we surveyed self-declared disease ecologists and conducted a literature synthesis with machine-learning topic detection (Bird, Klein, & Loper, 2009; Blei, 2012; Loper & Bird, 2002). Systematic and quantitative approaches to literature syntheses are increasingly favored over narrative-based reviews (Haddaway & Watson, 2016; Hedges & Olkin, 2014; Lajeunesse, 2010). However, high volumes of published research make theme synthesis very difficult and require innovative approaches (Lajeunesse, 2016; Nunez-Mir, Iannone, Pijanowski, Kong, & Fei, 2016). Following recent adoptions of data mining approaches to systematic reviews (Han & Ostfeld, 2019), we apply topic detection using non-negative matrix factorization to characterize the research core and trajectory of disease ecology. More broadly, our approach can provide a quantitative synthesis framework to examine the frequency that topics are published in other fields that cross traditional disciplinary boundaries.

## MATERIALS AND METHODS

### Survey

We developed a survey questionnaire to quantify the demographics and research core of disease ecology (Pennsylvania State University Institutional Review Board Study 00010582; Supporting Information). The survey was disseminated on disease ecology email listservs such as conference attendees (e.g., the past five years of Ecology and Evolution of Infectious Disease conferences, Ecological Society of America), scientific organizations and networks (e.g., VectorBiTE, American Society of Parasitologists, Ecological Society of America disease ecology section), and institutional research centers (e.g., Pennsylvania State University Center for Infectious Disease Dynamics, University of Georgia Center for the Ecology of Infectious Diseases). We also distributed the survey to prominent non-USA research groups (in e.g., South America, Europe, Australia) to diversify our survey participants. However, we acknowledge that survey reach was heavily biased towards established research centers and active researchers in the disciplinary community, primarily in North America and Europe, and surely missed certain individuals and groups, particularly those who conduct relevant research on infectious disease but may not necessarily identify as disease ecologists (e.g., medical entomologists and historians).

The survey was open from November 2018 until January 2019, closing once the response rate dropped below two new responses per day for one consecutive week. All survey participants were self-declared disease ecology practitioners, who were informed about the potential use of results in a consent statement. The survey asked participants questions on their demographics, institution, education, types and topics of current research, and influential scientific articles and journals. It included a combination of multiple choice and short answer response questions. A full copy of the survey and description of the data cleaning procedure is available in the Supporting Information.

### Literature search

Our objective was to compile an extensive corpus robustly representative of publications in disease ecology rather than to include every article per se. Literature search terms are often generated by the authors, which may impose bias. To generate a list of search terms with reduced author-bias, we compiled a set of papers that cited the foundational paper in disease ecology and was considered to be highly influential to the survey participants (Anderson & May, 1979; Table 1). Using this set of papers, we performed topic detection algorithms (*nltk* library, Python 2.7; Bird et al., 2009) to generate the list of 13 base keywords (e.g., the word parasite could have multiple prefixes and suffixes) that were used to search the wider literature (see Supporting Information Literature Search Methods). To this end, our literature search terms emerged from disease ecology literature itself, and then were refined through the process described below and in Figure 1.

**Table 1.**
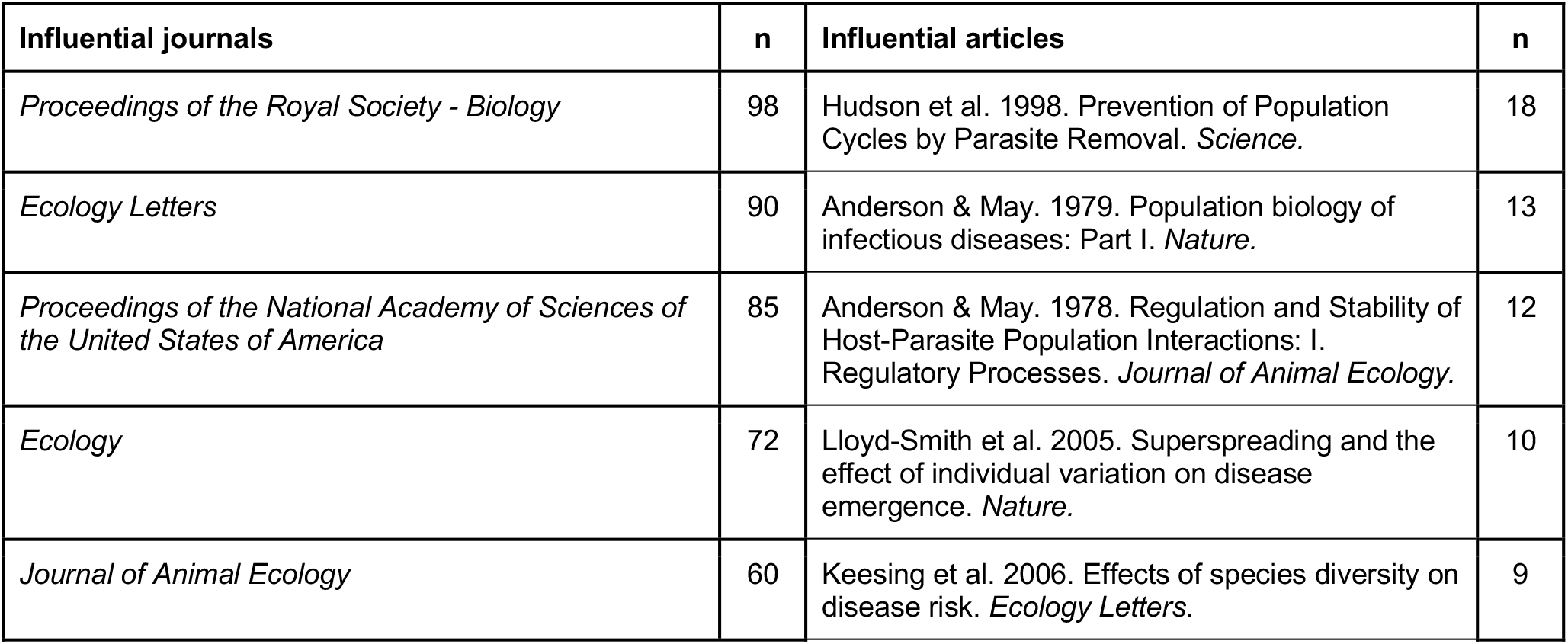
The five highest ranking scientific journals (left) and articles (right) based on survey responses. Survey participants were asked to pick journals other than *Science* or *Nature*.

**Figure 1.**
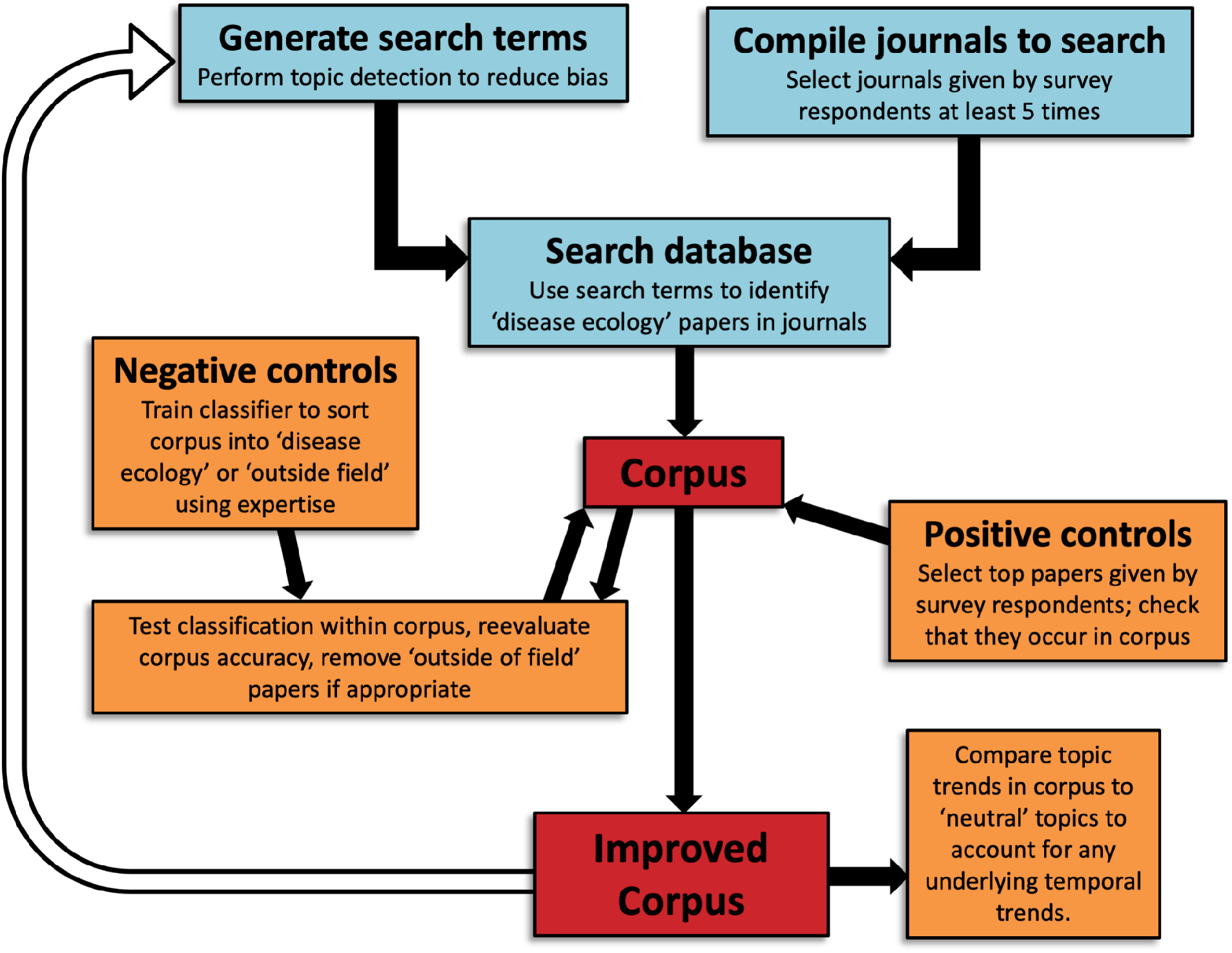
Workflow of systematic literature search and corpus development. Box coloration denotes different stages of corpus development: literature compilation (blue), corpus assessment and validation (orange), and corpus completion (red). The unfilled arrow denotes repeating of the workflow to optimize corpus accuracy.

**Figure 2.**
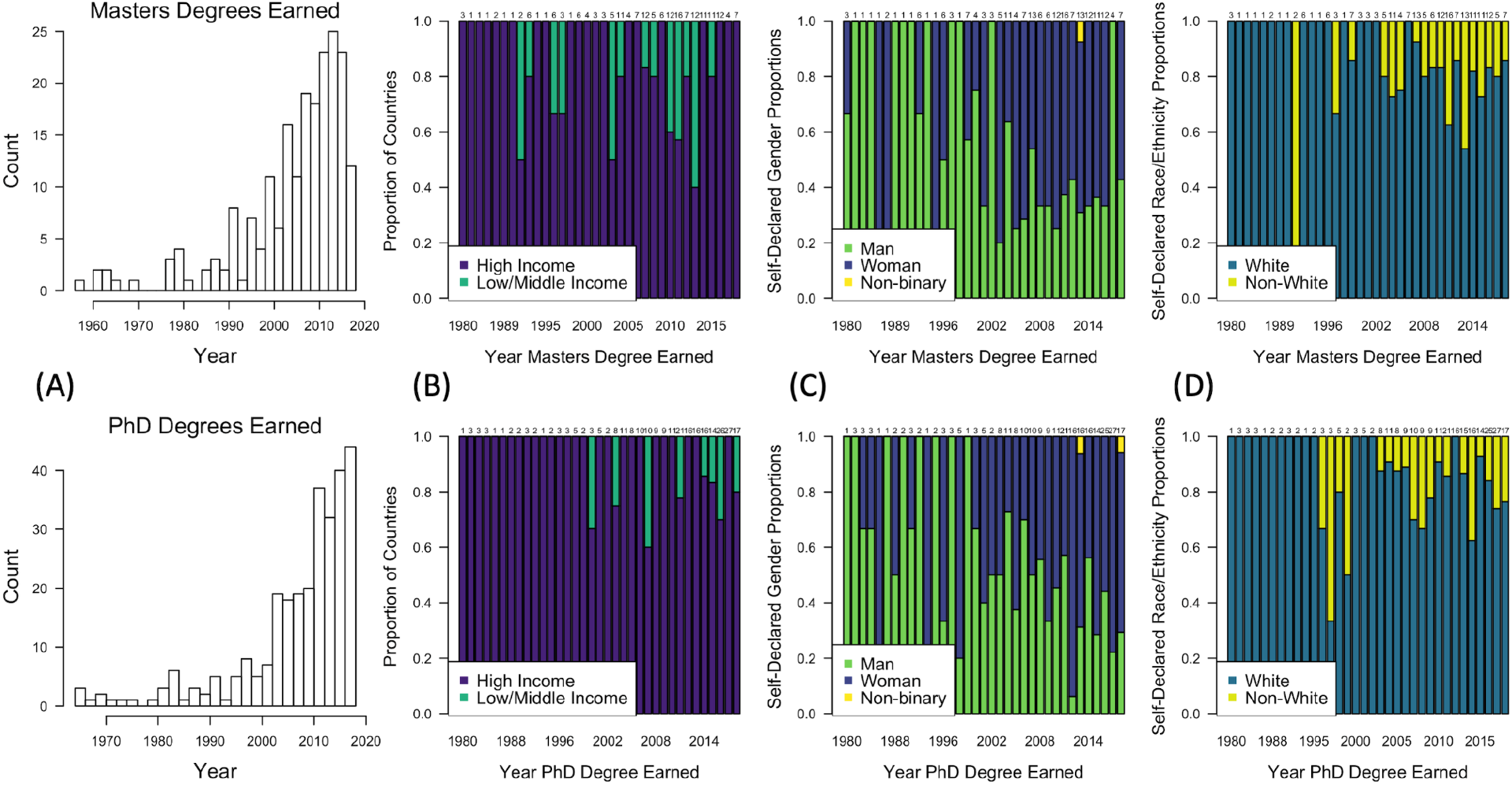
Plots showing demographic trends in survey participants that indicated they earned a Master’s (top row) or PhD (bottom row) degree by 2018. **(A)** Displays the number of degrees earned by year, and the stacked bar plots show the proportion of degree earners by **(B)** high or low/middle income countries, **(C)** self-declared gender identity, and **(D)** self-declared race/ethnicity. Sample sizes are displayed above their respective bars and years included 1980-2018; some years may be excluded (from the x-axis) if no degree earners indicated the information of interest.

**Figure 3.**
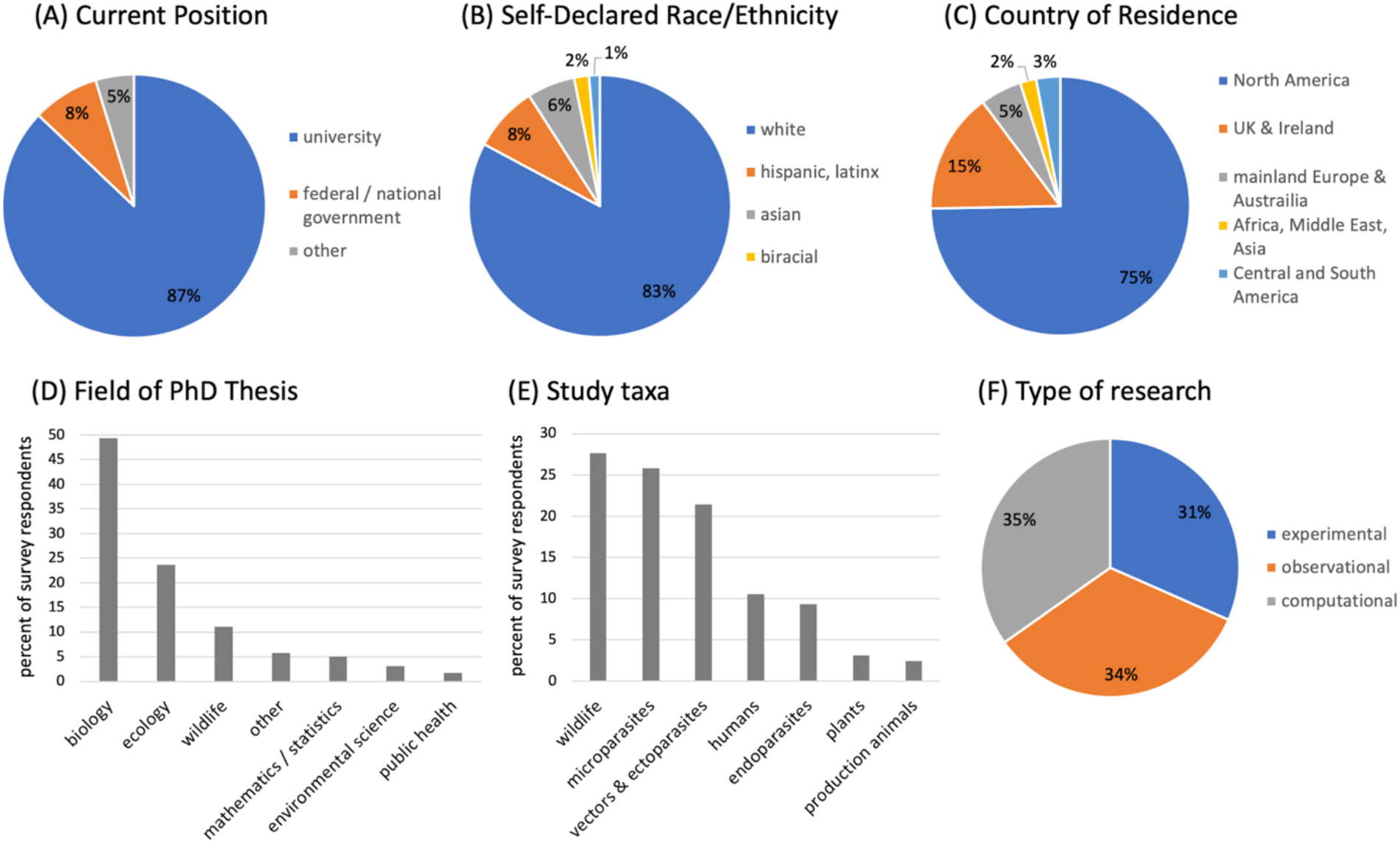
Summary of disease ecology demographics and research topics/types from survey participants. Pie charts display **(A)** current position or institution type, **(B)** self-declared race or ethnicity, and **(C)** country of residence; research topics and types are categorized by **(D)** field of PhD thesis, **(E)** current study taxa, and **(F)** type of primary research.

The final literature search was conducted in Web of Science for the years 1975 to 2018. Each article had to meet specific criteria using Boolean filters, including a focus on studying a pathogen or parasite, host infections (to distinguish from solely environmental persistence of microorganisms), and individual-level or higher-order dynamics (e.g., not cellular processes, with the exception of those analyzed as a population-level process). The full list of search terms is provided in the Supporting Information, alongside a set of exclusionary terms to remove similar but non-disease ecology articles. Web of Science categories were used to narrow our search and also reduce false-positive inclusions. To reduce bias, both search terms and included journals were based on survey results. We included journals that were listed by at least four survey participants as significant to the field (n = 42), as well as *Nature* and *Science*. Finally, articles with fewer than four citations were removed as a form of quality control.

To evaluate false-positives, two authors (DJB and KMF) independently evaluated the same 100 randomly selected articles and classified them as ‘disease ecology’ or ‘outside the field’. Papers that fell outside the field predominantly described pathogenesis, bacterial communities, or genetics/genomics (Fig. S5). Within-host studies were accepted if they focused on population-level processes (e.g., Cressler, Nelson, Day, & Mccauley, 2014) or parasite manipulation. 75% of the articles in the final corpus were classified as disease ecology, and consensus was strong among evaluators (94% agreement, Cohen’s κ=0.84). Within false-positive papers, there was no association between topical and temporal trends (χ^2^ = 72.84, p = 0.29, Supporting Information False-Positive Literature Assessment).

To evaluate false-negatives, we cross-validated our corpus using our survey data. Specifically, we assessed whether articles that were identified by at least two survey participants as influential were present in our corpus. We calculated the proportion of papers that were included in our corpus out of the list of such articles, with the requirement that at least 70% of papers had to be included. Of the influential articles identified by survey participants (written ≥2 times) restricted to journals used in building the corpus, approximately 71% (50/70) were present in the corpus. The ‘most influential’ articles had a higher probability of being included: the corpus included 85% of articles written four or more times, 75% of articles written three times, and 63% of articles written twice. We adjusted the search and exclusion terms twice using the workflow described in Figure 1 (unfilled arrow) to obtain a corpus with high classification and cross-validation success.

### Literature analysis

We conducted topic detection on the validated corpus using non-negative matrix factorization. Topic clusters represent a set of co-occurring words that can be used to define an area of research. The number of topic clusters (i) and words per topic (j) were the only parameters imposed on the literature analysis. To select appropriate values for i and j, we ran topic detection for a range of values and combinations of i and j and assessed outputs. If i was too small or too large, we were unable to detect temporal variation in that topic. If j was too small or too large, the topics were not clearly defined. For example, a topic with only five words may not be interpretable; similarly, a topic with 30 words may be too broad to assign meaning. We used I = 15 and j = 15, so our corpus was analyzed for 15 topics with 15 words each.

We used K-means clustering from the *nltk* Python library to construct topics, where each topic comprised 15 commonly co-occurring words. We assigned a name to each topic to describe its theme. For example, we named a topic containing immunodeficiency, HIV, patient, therapy, drug, AIDS, background, treatment, and risk, as an HIV topic. We gave each topic name a ‘confidence’ measurement of 1–3, from high to low confidence in identifying the topic (Supporting Information Topic detection). In addition to topics that emerged from the literature, we also generated and assessed our own topic lists based on key research areas: climate change, dilution effect, superspreaders, network analysis, EIDs, infectious diseases in bats and rodents, chytrid fungus, theoretical modeling, and field experiments (Fig. 4; full topic lists are in the Supporting Information). To ensure topic trends were not confounded by an increase in the total number of published articles through time, we constructed a baseline topic using neutral words that should be in all disease ecology articles: analysis, study, and paper. We evaluated temporal trends in publications for each theme using generalized additive models (GAMs) fit using the *mgcv* package in R (Wood, 2006). The proportion of words in each topic relative to all words was modeled as a binomial response using thin plate splines with shrinkage for publication year. Lastly, to assess covariation among topics, we estimated Spearman’s rank correlation coefficients (ρ) at the zero-year lag.

**Figure 4.**
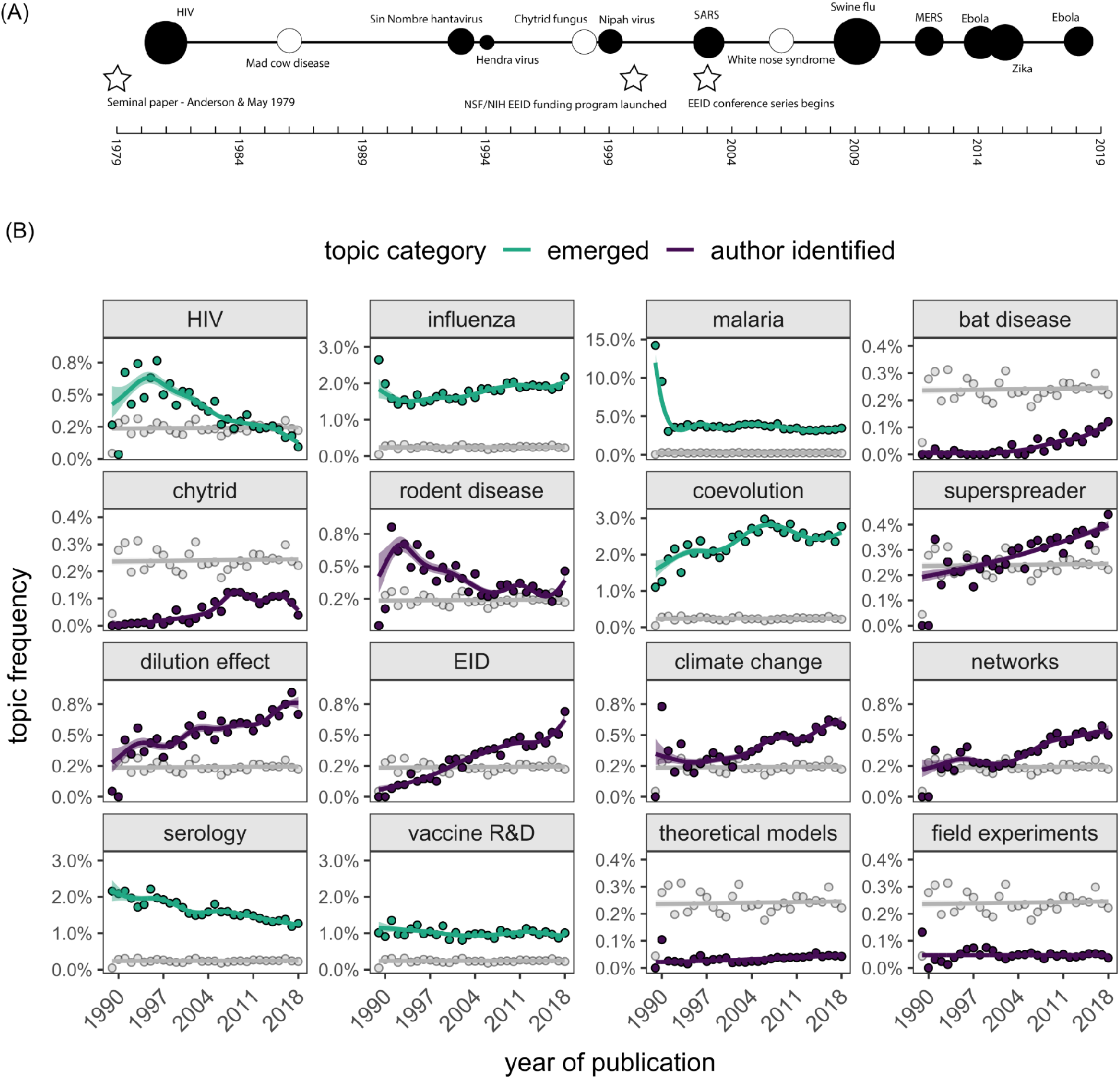
Timelines of events relevant to disease ecology and research trends. **(A)** A timeline of select human (filled circles) and wildlife (open circles) infectious disease events from 1979-2018. The approximate number of humans infected is represented by circle size (scaled by log/4). Analogous estimates are rare for wildlife diseases, thus these circles are an equivalent arbitrary size. Stars denote notable events for the development of the field of disease ecology. **(B)** Frequency of publication on selected topics (green = emerged from the literature corpus, purple = selected by authors) compared with the neutral topic (gray) from 1976-2018; 1976-1989 are binned in the first data point. Thick lines and ribbons show the fitted values and 95% confidence intervals from the GAMs. Plots are ordered by thematic categorization: host– pathogen study systems, concepts, and research methods/approaches.

## RESULTS

### Survey

A total of 413 self-declared disease ecologists participated in the survey. The average respondent was 36.1 years old (range: 21-76, median: 34, n=348; categorized as: ≤25, 26-30, 31-35, 36-40, 41-50, 51-60, >60). 76.7% of participants (n=408) considered at least half of their research to fall within disease ecology. Participants that considered ≥75% of their research to be disease ecology were concentrated from ages 26–40, and most self-identified as women (60%, n=344). More broadly, 56.1% of participants identified as women (n=231), 42.5% as men (n=175), 0.7% as non-binary (n=3), and 0.7% preferred not to say (n=3). We report on participants that chose to disclose a gender identity for results regarding gender.

Most participants identifying as women were younger (age ≤35) than most participants identifying as men (age 26–50). The youngest age category (≤25 years) was 68.9% women (n=45), and the oldest age category (>60 years) was 85.0% men (n=20). Current positions held by survey participants were: undergraduate student (1.2%, n=5), Master’s student (2.9%, n=12), PhD student (24.5%, n=100), post-doctoral researcher (21.1%, n=86), faculty (39.5%, n=161), researcher (9.1%, n=37), and other (1.7%, n=7). Participants identifying as women comprised most of each academic position except Master’s student and faculty (Table S1). In general, most PhD students and post-doctoral researchers were young and identified as women. Most Masters’ students were young and identified as men, and most faculty were middle-aged and identified as men (Tables S1–S3). Non-binary participants were distributed across age (<50) and position categories (n=3), as were participants who preferred not to provide a gender identity (n=3).

We identified clear trends in education and demographics of survey participants. 92.7% of participants had completed their undergraduate degree by 2018 (n=382), 50.0% had completed their Master’s degree by 2018 (n=206), and 73.3% had completed their PhD by 2018 (n=302). 8.3% of participants had also earned an additional degree, most commonly as a Doctor of Veterinary Medicine (n=18). Nearly half of participants had a PhD but not a Master’s degree (45.4%; n=187). The total number of graduate degrees earned per year among disease ecology researchers has dramatically increased since the year 2000 (Fig. 2A). Broadly, 50% of PhDs earned were in biology graduate programs (e.g., biology, biological sciences, microbiology), 25% were in ecology graduate programs (e.g., ecology, ecology and evolution, plant science), 11% were in wildlife or fisheries graduate programs (e.g., wildlife, fisheries, zoology, entomology, animal science), and <10% were in mathematics/statistics, environmental science, public health, or agricultural programs. Among surveyed disease ecologists from 1990-2018, biology programs have consistently comprised about half of all earned PhDs, with ecology closely tracking but slightly decreasing since 2005 (Fig. S2). Over the same time, PhDs in mathematics/statistics and wildlife/fisheries programs have slightly declined and remained at approximately 10% (Fig. S2).

Most participants identifying as non-white were less than 36 years old, indicating a recent, though minor, diversification of disease ecology practitioners. This was especially pronounced when analyzed by education: the average proportion of participants identifying as non-white who have earned a Master’s degree has nearly doubled from 1980-1999 to 2000-2018 (9.6% to 19.1%), and had risen to 23.9% in the last decade (Fig. 2D). The proportion of participants identifying as non-white who have earned a PhD slightly increased from 1980-1999 to 2000-2018 but has since remained stable (15-19%; Fig. 2D). The proportion of Master’s and PhD degrees earned in low- and middle-income countries has also recently increased (Fig. 2B).

The proportion of participants identifying as women who have earned Master’s degrees approximately doubled from a mean of 27.5% during 1980-1999 to 53.2% during 2000-2018. From 2000-2018, this proportion was approximately stable at around 55%. Similarly, the proportion of participants identifying as women who have earned PhDs substantially increased from a mean of 34.3% during 1980-1999, to 52.4% from 2000-2018; this proportion has continued to increase to 62.6% since 2010 (Fig. 2C).

The most popular areas of research within disease ecology were ranked as: epidemiology, mathematical modeling, population ecology, wildlife ecology/management, parasitology, community ecology, and infectious disease evolution/life history (up to 5 responses were included per person; n participants=410, n responses=1739). The least common areas included behavioral ecology, bioinformatics, field and laboratory techniques, movement ecology, virology, landscape ecology, and zoology.

Survey participants were asked to write in scientific journals and articles that they believed were the most influential in disease ecology (Table 1). Influential articles (written in at least twice, n=76) were most often published in *Scienc*e (n=17), *Nature* (n=10), *Trends in Ecology & Evolution* (n=9), *Ecology Letters* (n=8), and *Proceedings of the National Academy of Sciences* (n=6). Interestingly, however, *Proceedings of the Royal Society - Biology* was ranked as the most influential journal by survey participants, followed by *Ecology Letters, Proceedings of the National Academy of Sciences, Ecology*, and *Journal of Animal Ecology* (Table 1). See Supporting Information for full lists of both articles and journals.

### Literature search & analyses

We compiled a list of 42 journals that at least four survey participants said were the most important in disease ecology, plus *Science* and *Nature*. We searched these 44 journals for relevant articles in the field using the algorithm described above, and our final corpus comprised 18,695 articles. Our validation processes demonstrated that at least 75% of these articles were properly classified, and we did not detect any systematic bias in falsely-positive articles. Articles span from 1975 to 2018, with most published after 2000, indicating a rapid and considerable expansion of the field since early foundational work in population biology and vector-borne disease (Anderson & May, 1979; Hess & Hayes, 1970; May & Anderson, 1979). However, some journals were not available in Web of Science until the 1980s-1990s, so article availability may slightly bias our corpus in the early years.

Topic clusters were classified into two categories: (1) those that emerged and were identifiable from the literature and (2) those that we deliberately searched for using key term searches. Of emergent topics, malaria and mosquito-borne pathogens appeared most frequently in the topic clusters (3/15), followed by experimental infection trials (2/15). Other clear topics included HIV, influenza, vaccine research, and host-pathogen coevolution. Some topics were more ambiguous but still identifiable, such as wildlife pathogens, tick-borne pathogens with rodent hosts, and serological analyses. Overall, we had high confidence assigning names to topic clusters emerging from the literature, indicating defined areas of research in the corpus (see Supporting Information).

Many of the topics that emerged from the disease ecology literature, such as malaria, influenza, and vaccination research and development, have remained constant in publication frequency over time (Fig. 4B). Others, such as HIV and serology, have declined in publication frequency over time, and host-pathogen coevolution has instead steadily increased. These emergent topics comprised a notable portion of the disease ecology literature and were more prominent in the literature than author-selected topics. A neutral topic, constructed for comparison, had a constant publication frequency through time (Fig. 4B, gray line in panels), thus validating the observed temporal changes in these topics.

Using key term searches, we next explored the frequency of publication of select topics: climate change, emerging infectious diseases (EIDs), the dilution effect, superspreaders, network analysis, pathogens in rodents and bats, chytrid fungus in amphibians, theoretical modeling, and field experiments (Fig. 4B). As with emergent topics, our topic detection was sensitive to detecting changes in frequency over time, identifying peaks and troughs. For instance, published research on pathogens in bats had a small peak around the time of the first SARS epidemic (2002-2004), and bat disease research has steadily increased since 1979; however, overall, literature on bat pathogens has been a small proportion of all disease ecology literature. Published research on rodent pathogens was greatest in the 1990s and has generally declined, although recent years have also seen an increase in rodent-related disease ecology publications. The frequency of publications on many topics has steadily grown and will likely continue to grow based on this trend, such as EIDs, climate change, the dilution effect, network analyses, and superspreaders. Chytrid fungus literature, on the other hand, appears to have declined or plateaued in recent years. Lastly, publication frequency of both theoretical modeling of infectious disease and field experiments has remained constant but rare over time.

Frequencies of published topics displayed strong degrees of cross-correlation, with both positive and negative covariation in annual trends (Spearman’s rank correlation coefficient ρ=-0.89-0.88; Fig. S8). Particularly strong positive correlations were observed for superspreaders and network analyses (ρ=0.88), superspreaders and the dilution effect (ρ=0.85), EIDs and bats (ρ=0.83), EIDs and the dilution effect (ρ=0.83), EIDs and network analysis (ρ=0.82), and superspreaders and bat pathogens (ρ=0.80). Especially strong negative correlations were observed for EIDs and serology (ρ=-0.89), serology and the dilution effect (ρ=-0.87), serology and bat disease (ρ=- 0.81), influenza and HIV (ρ=-0.77), climate change and HIV (ρ=-0.76), network analysis and serology (ρ=-0.70), influenza and rodent disease (ρ=-0.68), climate change and serology (ρ=- 0.68), malaria and network analysis (ρ=-0.65), and malaria and rodent disease (ρ=-0.65).

## DISCUSSION

Interdisciplinary research fields can rapidly grow to address important societal needs, and retrospective analysis of their evolution can help improve their future trajectory and growth. By combining a survey with a powerful quantitative literature synthesis, we here demonstrate the increasing gender and institutional diversity of disease ecology practitioners alongside the breadth of research activities. Certain topical themes that emerged from our literature corpus, such as influenza, malaria, and vaccine research and development, have remained prominent foci of disease ecology, whereas an increase in the frequency of publication of a priori selected topics such as emerging infectious diseases, climate change, and effects of biodiversity loss emphasize how this expanding field has mirrored global events and priorities.

Self-declared disease ecology practitioners are becoming more diverse in terms of country of education, gender identity, and institution (Fig. 1). The gender trends identified here are echoed in engineering, computer science, and mathematics/statistics where the proportion of women earning graduate degrees has increased over the past two decades (20-43% of Master’s and Doctorates earned in 2014), yet remains low in physics (18.7% of Doctorates earned in 2014) (National Science Foundation, 2020). Women authorship has increased in ecology and evolution literature every year from 2009-2015 (Fox, Ritchey, & Paine, 2018), and the proportion of women journal editors also increased over that time but was still low relative to men (Fox, Duffy, Fairbairn, & Meyer, 2019). In terms of race/ethnicity, the rate of people identifying as Hispanic earning Bachelor’s degrees in science and engineering has increased slowly since the 1990s, but remained approximately constant for people identifying as black, African American, or Asian (National Science Foundation, 2020).

Diversity in the workplace and educational institutions is fundamentally important and increases performance, cooperation, problem-solving, and student retention (Drury, Siy, & Cheryan, 2011; Milem, 2003; Roberge & van Dick, 2010). The highest demographic and institutional diversity we identified was in younger age groups (<36 years old), graduate programs, and post-doctoral positions (Table S2, S3). This may be due to increasing levels of education globally (Group of Eight, 2013; UNESCO Institute for Statistics, 2020), or targeted programs to increase diversity in science and mathematics, particularly focused on recruiting women (Burke & Mattis, 2007; Huntoon & Lane, 2007). Another non-mutually exclusive driver of these trends could be the failed retention of minorities and women in later career stages (Blickenstaff, 2005; Diekman, Brown, Johnston, & Clark, 2010; Shaw & Stanton, 2012). Yet importantly, although we identified some relative increases in diversity within disease ecology, the field as a whole remains quite homogenous in terms of gender, race and ethnicity, and geography, and marginalized groups face considerable inequities and discrimination in science fields – e.g., harassment and exclusion, less likely to have grants funded, hold fewer faculty positions, and have more limited access to academic experiences and resources (Allen, Epps, Guillory, Suh, & Bonous-Hammarth, 2000; M. S. Jones & Solomon, 2019; Rissler, Hale, Joffe, & Caruso, 2020). Concerted efforts to improve equity must continue and explicitly address recruitment and retention, especially for fostering racial and ethnic diversity.

We acknowledge that surveys can be a biased source of information because researchers rely on voluntary participation. For example, studies of academic survey participation have shown that people who identify as women are more likely to respond to surveys, while academic rank had little influence on response rate (e.g., tenured versus tenure-track) (Saleh & Bista, 2017; Smith, 2008). Additionally, as our survey was shared via email listservs, it is likely that many people did not see or receive our request for participation. Because the field of disease ecology is relatively new and multidisciplinary, it is more challenging to identify smaller research groups both at universities (i.e., individual lab groups) or decentralized working groups (e.g., Bat One Health Research Network (BOHRN)). Our survey dissemination and participation likely reflects broader geographic biases in ecology research and publishing (Nuñez, Chiuffo, Pauchard, & Zenni, 2021), which could subsequently affect the influential literature identified (Table 1); however, we were unable to assess these limitations. While there are shortcomings of surveys, they remain a widely used method of data collection, and the survey developed here provides the first description of the composition of disease ecologists and important literature that we hope is built upon in the future.

The second part of our study comprised an extensive literature synthesis. Literature reviews can be compromised by author bias when search terms are subjectively selected (reviewed in Okoli, 2015). There is a trade-off between the scope and errors when constructing a literature corpus. For example, narrower ecological literature reviews usually consist of a <2,000 article corpus and often much fewer (Han & Ostfeld, 2019; Lowry et al., 2013; Poff & Zimmerman, 2010; Wortley, Hero, & Howes, 2013), and papers may be individually-assessed for inclusion (e.g., Uehlinger, Johnston, Bollinger, & Waldner, 2016) sample size = 9 papers). Our analysis, on the other hand, captured a diverse range of literature topics within a broader field, resulting in a corpus of over 18,500 papers. We quantified the false-positives (type I error) and true-positives in our corpus, which is rarely accounted for or reported in ecological literature reviews (Haddaway & Watson, 2016). False-positives are inevitable in such a large body of literature, but the false-positive papers identified were unbiased with respect to year or topic. Our true-positive rate was high – 85% of articles written in by participants four or more times were present in our corpus. Therefore, using our robust corpus formulation and validation (Fig. 1), we are confident that we were able to identify true, broad patterns across the disease ecology literature at a large scale.

Topic detection of publications revealed how published research priorities changed over time. For example, the frequency of published HIV research peaked in the 1990s but in recent years has declined to be lower than the frequency of most other topics in the field. While a direct association is difficult to demonstrate, the decrease in HIV-related publications has roughly coincided with advances in HIV treatments (hiv.gov, 2019). Likewise, although we also observed significant temporal fluctuations in publications related to malaria, their frequency has remained remarkably constant since the late 1990s. This likely reflects ongoing and continued efforts to reduce public health burdens of this disease and to understand complex interactions between mosquito vectors, human hosts, and the environment (Suh et al., 2020), especially in the context of emerging human pathogens (e.g., *Plasmodium know*lesi, Lee et al., 2011). We also identified broader concept-based trends in disease ecology literature. In particular, the frequency of published research on the dilution effect has undergone several spikes following key findings (Civitello et al., 2015; Keesing, Holt, & Ostfeld, 2006) and a steady increase in publication rate. Similarly, predicting how climate changes may alter pathogen spread continues to be of growing research interest (Altizer et al., 2013; Ryan, Carlson, Mordecai, & Johnson, 2019), as does research on superspreaders (Lloyd-Smith, Schreiber, Kopp, & Getz, 2005). More broadly, these temporal patterns in publications suggest that work related to biodiversity and infectious disease, climate change, pathogen spillover, heterogeneity in pathogen transmission, and new tools to analyze epidemiological data will all continue to be active areas of inquiry.

Published research on emerging infectious diseases and bats has increased through time, consistent with bats being established reservoir hosts for pathogens such as Nipah virus, SARS, Marburg virus, and Hendra virus, and with infection-related population declines in bats through white-nose syndrome (Calisher, Childs, Field, Holmes, & Schountz, 2006; Frick et al., 2010; Fig. 4A). However, although the frequency of disease ecology publications on bat pathogens have increased markedly in recent years, they still remain relatively understudied compared to our neutral term and rodent pathogens, and comprise only a small proportion of emerging infectious disease research in general. It is worth noting that our search does not include the SARS-CoV-2 pandemic, which we expect to lead to a large spike in disease ecology research on pathogens with their evolutionary origins in bats and the role of intermediate hosts in spillover (e.g., Andersen et al., 2020).

In general, published research on epidemics tended to lag rather than precede events, such that we observed a spike in the frequency of publications on high-profile pathogens followed by a decline or plateau (e.g., chytrid fungus). Emergent topics were remarkably stable through time, with the exception of HIV and host-pathogen coevolution, which have respectively decreased and increased. The frequency of published research focusing on concepts (e.g., the dilution effect, superspreaders, coevolution) or approaches (e.g., network analyses) rather than specific hosts or pathogens tended to rise more gradually and remain a notable proportion of the literature. On the other hand, mosquito-borne pathogens and influenza have been defining topics over the entire time series, which we expect to persist for the foreseeable future. We observed exceptions for theoretical and field experimental approaches to disease ecology; on the one hand, publications with these approaches have remained constant over time. However, publication frequency was rare relative to the broader disease ecology literature. This could signal that these approaches are relatively rare, but we suspect that publications using theoretical modeling or field experiments may not use the same set of co-occurring words, thus making them harder to identify as distinct using topic detection.

Although our analysis of cross-correlation between the topic frequency time series is associative, we observed several interesting relationships. The frequency of publications on bat disease, chytrid fungus, climate change, the dilution effect, superspreaders, and emerging infectious diseases were all positively correlated over time, suggesting a general increase in research on emergent disease risks to wildlife and humans in relation to anthropogenic change and heterogeneities in pathogen transmission (Daszak et al., 2001; K. E. Jones et al., 2008; Lloyd-Smith et al., 2005). From another perspective, the frequency of publications on serology have steadily declined over time and was negatively associated with topics such as bat disease and emerging infectious diseases, possibly indicating advances in rapid and affordable sequencing efforts to quantify pathogen diversity (Lipkin, 2013). These associations provide testable hypotheses for a future analysis that examines the combination of concepts and methods in published literature.

Using a survey and quantitative literature synthesis, this study demonstrates that disease ecology is a rapidly growing field, albeit one that will require continued efforts to enhance recruitment and retention to improve diversity. Our analysis identified trends and publication patterns of research topics addressed by disease ecologists. More broadly, our quantitative synthesis framework could help examine the composition and trends of other major research topics that cross traditional disciplinary boundaries.

## Supporting information

SupportingInformation

## Acknowledgements

Our survey was considered exempt under the Pennsylvania State University Institutional Review Board (STUDY00010582). We thank all participants who completed the online survey as well as helpful feedback from participants at the 2019 Ecology and Evolution of Infectious Diseases conference. We are grateful to Vanessa Ezenwa and Peter Hudson for constructive comments on an earlier version of this report.

