## SupportingInformation for "Demography, education, and research trends in the interdisciplinary field of disease ecology"

**Survey**
Demographic, institution, and education questions were write-in response or allowed participants to select from a set of choices. Research questions asked participants to select from a set of choices, allowing one or multiple selections. Respondents were asked to provide self-identifying gender identity and race/ethnicity from category options (US Census Bureau accessed 2019), but with the option to include additional information if desired. The survey did not require any questions to be answered before moving on. We decided which topics to include as choices by brainstorming with colleagues and experts in disease ecology, as well as considering the breadth of topics in disease ecology publications. This survey was approved by the Pennsylvania State University Institutional Review Board Study 00010582.

***Email prompt***
*We are conducting a study to better understand the background and diversity of disease ecologists, and how this relatively new discipline has developed over time. We're asking anyone who identifies any part of their research as disease ecology, regardless of how much or how little, to please complete this short online survey (approx. 10 minutes). We're hoping to get strong representation from different career stages (e.g. professor, postdoc, grad student), so please encourage your colleagues to take it!*

*(Link here)*

*The success of this study is reliant on high participation. We're very grateful for your contribution, and look forward to sharing the study results with you in due course.*

*Kind regards from the study team,*
*Ellen Brandell, Kristian Forbes, Daniel Becker, and Laura Sampson*

***Consent statement***
Please note that this information is optional and for our records or broad summary statistics only, e.g. the proportion of respondents of each gender and career stage. This information will be stored securely and anything that could lead to personal identification will be removed from reports. By taking this survey, you are consenting that your non-identifiable responses may be published and publicly available.

***Survey dissemination***
Below is a list of the email groups that we shared the survey with:
- Ecology and Evolution of Infectious Disease conference attendees (i.e. those that registered and provided an email address) 2014-2019
- Pennsylvania State University Center for Infectious Disease Dynamics
- VectorBiTE
- Bozeman Disease Ecology Group
- Ecological Society of American Disease Ecology Section presenters 2014-2019
- American Society of Parasitologists
- British Ecological Society Parasite and Pathogen Special Interest Group

- University of Georgia Center for the Ecology of Infectious Diseases
- Mammal-I
- University of Helsinki Metapopulation Group
- Griffith Wildlife Disease Ecology Group
- University of Melbourne One Health Research Group
- University of Antwerp Evolutionary Ecology Group
- EDEN / EDENext consortiums

***Cleaning survey results***
Full surveys were removed from the data if there were duplicates (e.g. if the form was submitted multiple times by the same person; n = 1), or if respondents were retired (n=2). Otherwise, individual responses were only removed when instructions were not followed properly (e.g., publication information entered was incomplete and could not be deciphered) or questions were not answered; the total number of responses is indicated for each results section of the main text document. We corrected typos or responses that were entered incorrectly but decipherable. When relevant, responses were categorized to allow meaningful analysis. For example, current position was consolidated into: undergraduate student, Master's student, PhD student, post-doctoral researcher, faculty, scientist, and other. Similarly, institutions were grouped by country and degree departments/fields into categories (e.g., 'agriculture', 'poultry science', etc.; 'wildlife', 'fisheries', 'zoology', 'entomology', etc.; 'biology', 'biological sciences').

**Additional Survey Results**
We acknowledge potential sampling biases that arise from any voluntary survey. Firstly, we did not sample all disease ecologists. We were limited by distributing our survey to visible, participatory groups and may have missed smaller lab groups, especially outside of the USA and Europe. Nonetheless, we distributed our survey to thousands of people in disease ecology related groups and had global participation. Secondly, there is evidence that women are more likely to participate in voluntary surveys (Smith 2008), and gender identity and race can influence how survey questions are interpreted and answered (Rooney et al. 2005), both of which may bias survey results. Given the diversity of our dataset, however, we do not believe these were prevailing problems.

Here we provide responses for degrees earned or in progress through 2019, as numerous survey participants were working towards a degree and represent a portion of the field. However, in the main text, we only included individuals that had completed their degree (i.e., through year 2018) to ensure the data presented represent accurate values.

***Survey results – demographics, education, current position***
Of 402 participants that provided an institution or current position, 87.1% were from a university, 8.2% were from the federal government, and 4.7% fell into other categories (e.g., state government agencies, museums, research centers, industry). Most survey participants identified as white (81.4%; n=324), followed by hispanic or latinx (8.0%; n=32), asian (5.8%; n=23), biracial (1.8%; n=7), and black or African American (1.3%; n=5) (Fig. 3B).

**Table S1**. Proportion of each category by gender identity.

| | *female* | *male* | *other* |
|---|---|---|---|
| *Undergraduate student* | 80.0% (4) | 20.0% (1) | 0 |
| *Master's student* | 33.3% (4) | 58.3% (7) | 8.3% (1) |
| *PhD student* | 71.7% (71) | 28.3% (28) | 0 |
| *Post-doctoral researcher* | 60.0% (51) | 38.8% (33) | 1.2% (1) |
| *Faculty* | 47.5% (76) | 52.5% (84) | 0 |
| *Researcher* | 59.5% (22) | 40.5% (15) | 0 |
| *Other* | 28.6% (2) | 57.1% (4) | 14.3% (1) |

**Table S2**. Proportion of each age bin by gender identity.

| | *female* | *male* | *other* |
|---|---|---|---|
| <= *25* | 68.9% (31) | 28.9% (13) | 2.2% (1) |
| *26-30* | 60.5% (46) | 38.2% (29) | 1.3% (1) |
| *31-35* | 74.1% (60) | 25.9% (21) | 0 |
| *36-40* | 50.8% (30) | 49.2% (29) | 0 |
| *41-50* | 41.2% (21) | 56.9% (29) | 2.0% (1) |
| *51-60* | 50.0% (7) | 50.0% (7) | 0 |
| *60+* | 15.0% (3) | 85.0% (17) | 0 |

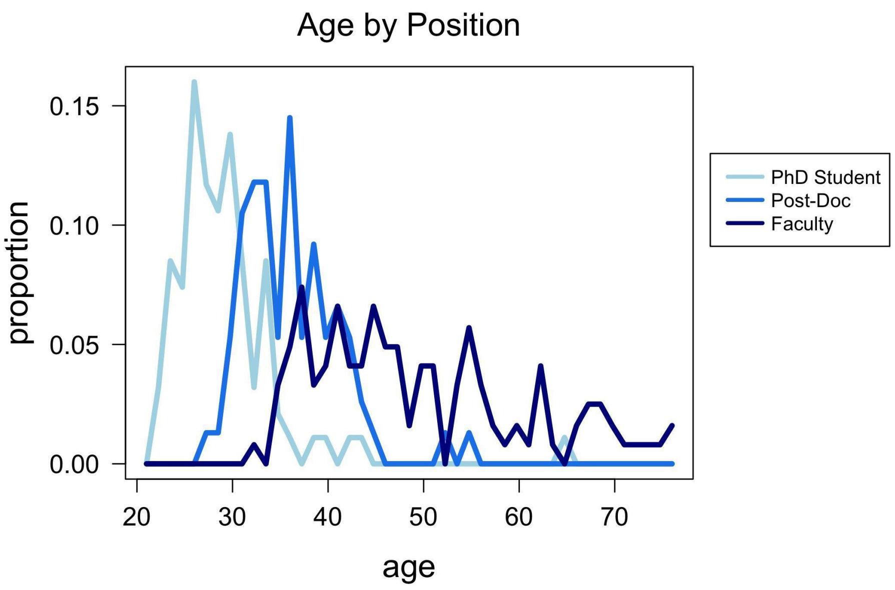


**Figure S1**. Proportion of PhD students (light blue), post-doctoral researchers (medium blue), and faculty (dark blue) within each year of age from 20 to 76 years old.

**Table S3**. Proportion of each age and position category by gender identity (n=343).

| **Undergraduate student** | *female* | *male* | *other* |
|---|---|---|---|
| <= *25* | 40.0 % (2) | 0 | 0 |
| *26-30* | 20.0% (1) | 20.0 (1) | 0 |
| *31-35* | 20.0% (1) | 0 | 0 |
| **Master's student** | | | |
| <= *25* | 16.7% (2) | 58.3% (7) | 8.3% (1) |
| *26-30* | 8.3% (1) | 0 | 0 |
| *31-35* | 8.3% (1) | 0 | 0 |
| **PhD student** | | | |
| <= *25* | 28.7% (27) | 6.4% (6) | 0 |
| *26-30* | 30.9% (29) | 17.0% (16) | 0 |
| *31-35* | 10.6% (10) | 2.1% (2) | 0 |
| *36-40* | 1.1% (1) | 2.1% (2) | 0 |
| *41-50* | 0 | 0 | 0 |
| *51-60* | 1.1% (1) | 0 | 0 |

| | | | |
|---|---|---|---|
| **Post-doctoral researcher** | | | |
| *26-30* | 17.1% (13) | 11.8% (9) | 1.3% (1) |
| *31-35* | 32.9% (25) | 13.2% (10) | 0 |
| *36-40* | 9.2% (7) | 11.8% (9) | 0 |
| *41-50* | 1.3% (1) | 1.3% (1) | 0 |
| **Faculty** | | | |
| *26-30* | 0 | 0.8% (1) | 0 |
| *31-35* | 13.1% (16) | 5.7% (7) | 0 |
| *36-40* | 13.9% (17) | 11.5% (14) | 0 |
| *41-50* | 13.9% (17) | 19.7% (24) | 0 |
| *51-60* | 4.1% (5) | 5.7% (7) | 0 |
| *60+* | 1.6% (2) | 9.8% (12) | 0 |
| **Researcher** | | | |
| *26-30* | 7.4% (2) | 3.7% (1) | 0 |
| *31-35* | 14.8% (4) | 7.4% (2) | 0 |
| *36-40* | 18.5% (5) | 14.8% (4) | 0 |
| *41-50* | 11.1% (3) | 11.1% (3) | 0 |
| *51-60* | 3.7% (1) | 0 | 0 |
| *60+* | 3.7% (1) | 3.7% (1) | 0 |
| **Other** | | | |
| *26-30* | 0 | 14.3% (1) | 0 |
| *31-35* | 28.6% (2) | 0 | 0 |
| *36-40* | 0 | 0 | 0 |
| *41-50* | 0 | 14.3% (1) | 14.3% (1) |
| *51-60* | 0 | 0 | 0 |
| *60+* | 0 | 28.6% (2) | 0 |

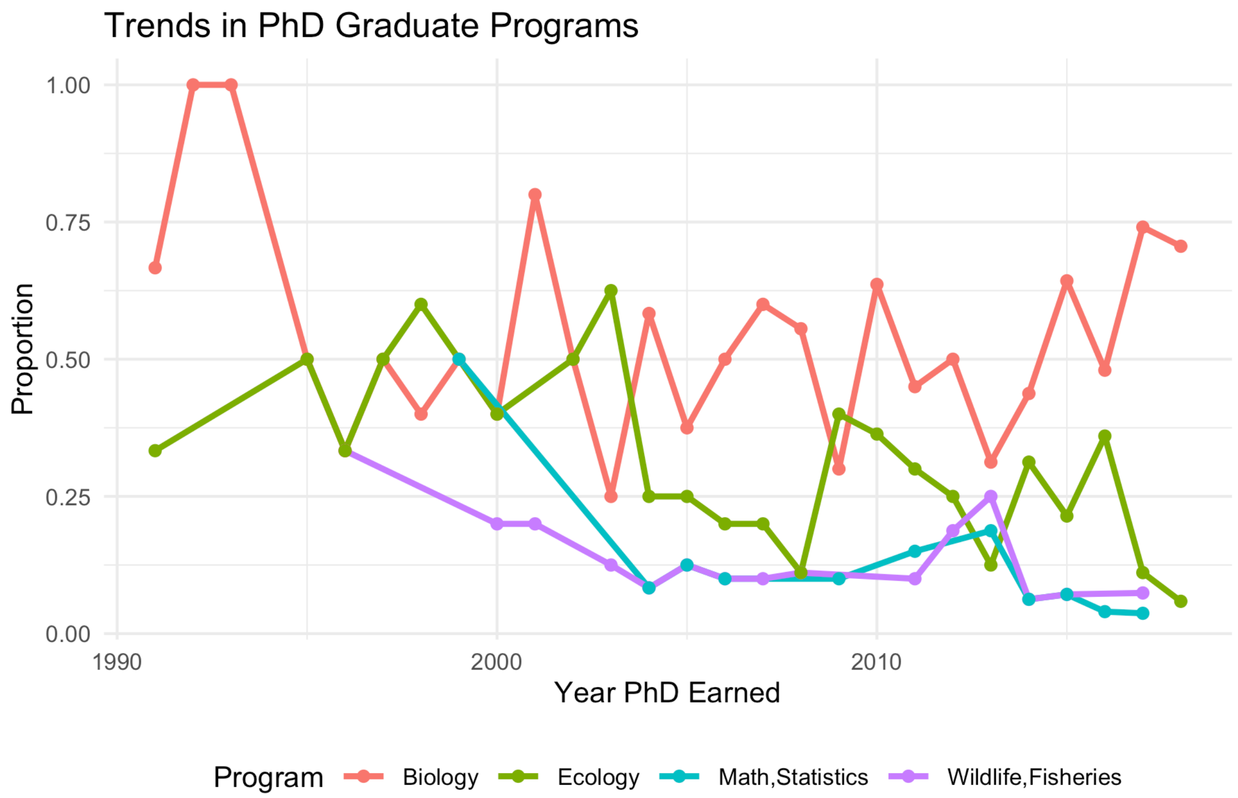


**Figure S2**. Proportion of PhDs earned in graduate programs from 1990-2018. Four broad categories describe 92% of survey participants: biology (biology, biological sciences, microbiology, or similar), ecology (ecology, ecology and evolution, plant science), math/statistics (mathematics, statistics, engineering, bioinformatics), and wildlife/fisheries (wildlife, fisheries, zoology, entomology, animal science). Annual sample sizes were small, ranging from 1 (1993) to 27 (2017), with an average of 9.7.

***Survey results – institution, current research, subdiscipline rankings***

When considering geographical location, 74.7% of participants reside in the USA or Canada (n=301); 15.1% reside in the UK/Ireland (n=61); 5.2% reside in other European countries or Australia (n=21); 2.0% reside in African, Middle Eastern, or Asian countries (n=8); and 3.0% reside in South or Central American countries (n=12) (Fig. 3C). Most PhDs earned were classified as biology (49.3%; n=177), followed by ecology (23.7%; n=85). Other degree departments/fields were much fewer: 11.1% wildlife (n=40), 5.8% other (n=21), 5.0% mathematics or statistics (n=18), 3.1% environmental science (n=11), 1.7% public health (n=6), and 0.28% agriculture (n=1) (Fig. 3D). Master's degree departments/fields were similar.

List of country of current institutions given by survey respondents (n=403):

| | | |
|---|---|---|
| Albania (1) | China (1) | Ghana (1) |
| Argentina (6) | Egypt (1) | Iran (1) |
| Australia (4) | Finland (5) | Ireland (3) |
| Belize (1) | France (1) | Italy (1) |
| Canada (12) | Germany (2) | Nigeria (1) |

Peru (1)
Poland (1)
Portugal (2)
Puerto Rico (1)
Qatar (1)
South Africa (1)
Spain (2)
Sweden (2)
Thailand (1)
UK (58)
USA (289)

Institution type by category (n=402, removed one reitree, Fig. 3A):
87.1% University (350)
8.2% Federal government (33)
4.7% Other (19)

We asked survey participants to indicate what proportion of their work or research fell into the field of disease ecology (n=408):
8.8% <25% (36)
14.5% 25-50% (59)
17.4% 51-75% (71)
28.9% 76-99% (118)
30.4% 100% (124)

We asked participants to record the two taxonomic groups they primarily study, but most participants recorded many. Therefore, results about research taxonomy include up to five responses per person. Some people wrote in a taxonomy classification, which we incorporated when it was coherent, otherwise it was removed. Here we give the proportion of each classification given – note that we combined ectoparasite and vector in the main text as we believe participants may have mistaken the two. Most participants study wildlife (27.6%; n=221), microparasites (25.8%; n=206), vectors/ectoparasites (21.4%; n=171), and humans (10.5%; n=84). The least studied taxonomic groups include endoparasites (9.3%; n=74), plants (3.1%; n=25), and production animals (2.4%; n=19) (Fig. 3E). The most common co-occurring pairs of taxonomic study groups were wildlife-microparasite (18.8%), ectoparasite/vector-wildlife (12.6%), and human-vector (9.6%). The least common co-occurring paris were: microparasite-vector (8.1%), wildlife-vector (7.2%), wildlife-ectoparasite (5.4%), wildlife-endoparasite (5.1%), and human-microparasite (4.7%)

Out of 400 participants, 23.75%, 28.50%, and 47.75% conducted applied, fundamental, or both applied and fundamental research, respectively. Approximately one-third of participants conducted experimental (31.6%), observational (33.6%), and computational research (34.8%) (n=351, Fig. 3F). Finally, participants predominantly conducted laboratory research (45.2%), then field research (39.7%) or both (15.1%) (n=398).

Survey respondents indicated their top five areas of research (n=410). Areas written in primarily incorporated into predetermined categories. The proportion of each research area are:
0.3% behavioral ecology (6)
2.0% bioinformatics (34)
5.7% biostatistics / empirical inference (99)
7.4% community ecology (128)
4.2% entomology (73)
13.2% epidemiology (230)
2.1% field techniques (36)
5.1% genetics / genomics (89)

3.0% immunology (52)
7.2% infectious disease evolution / life history (126)
2.0% laboratory techniques (35)
3.2% landscape ecology (55)
8.9% mathematical modeling (155)
2.5% microbiology (44)
2.6% movement ecology (46)
0.5% other (9)
7.5% parasitology (130)
8.3% population ecology (145)
2.8% virology (49)
8.1% wildlife ecology/management (140)
3.3% zoology (58)

The most common answers (≥100 respondents) were: epidemiology, mathematical modeling, population ecology, wildlife ecology/management, parasitology, community ecology, infectious disease evolution / life history (descending). The least common answers (<60 respondents): behavioral ecology (or similar write-ins), bioinformatics, field and laboratory techniques, movement ecology, virology, landscape ecology, zoology (descending).

We were interested in what areas of research disease ecologists considered to be the most important in the field. Therefore we asked respondents to rank given areas of research from 1 (most important) to 10 (least important) (n=152, Fig. S3). The most important area of research was, by far, ecology, then infectious disease evolution / life history, epidemiology, and field and laboratory techniques (descending). The least important areas of research were ecology, parasitology, immunology, field and laboratory techniques, microbiology/virology, and genetics/genomics/bioinformatics (ascending). It was interesting that ecology was ranked as both the most and least important area of research. Perhaps this was an artefact of incorrectly responding to this question (i.e., ranking backwards from 10 to 1 instead of 1 to 10), or perhaps this is indicative of a strong split in the field in regards to what disease ecologists deem as important research.

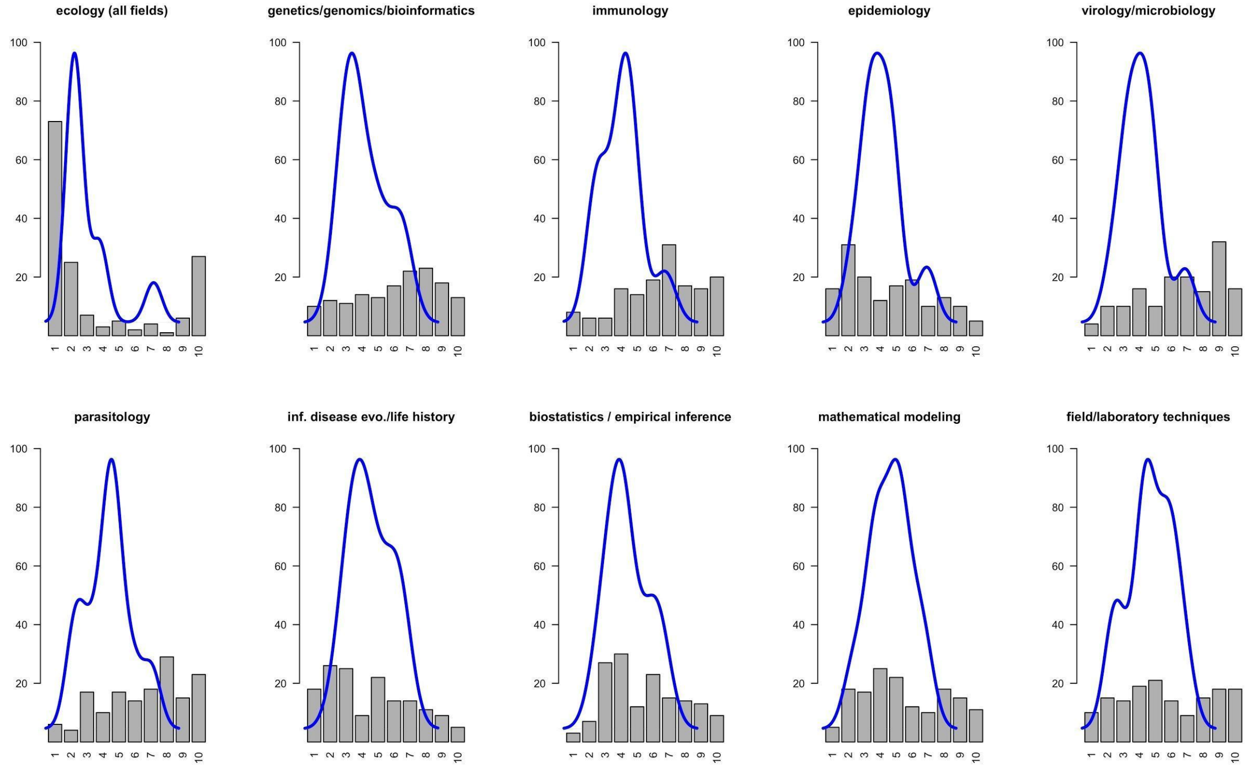


**Figure S3**. Density distributions overlaid on histograms for the ranks of each area of research. The ranks are displayed on the x-axis from 1 (most important) to 10 (least important); the y-axis is the frequency of responses.

Finally, we were interested in whether or not survey respondents were more likely to indicate that their personal areas of research were also the most important areas of research. We matched people's five selections for their areas of research with the top three areas they ranked as most important. Nearly half of participants believed two of their top areas of research were the most important (42.5%), only 15.7% believed three of their top research areas were the most important, and the rest believed their research areas were not the most important in disease ecology (Fig. S4). More simply, most people thought their own research fell into at least two of the three most important areas in the field.

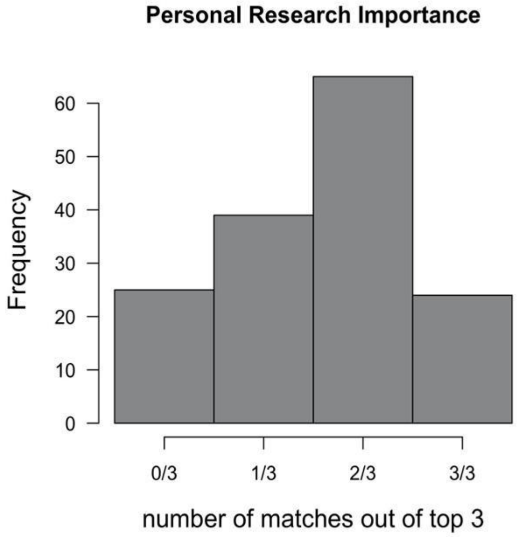


**Figure S4**. Histogram of how many personal areas of research were also the most important areas of research in disease ecology, according to survey respondents. The x-axis is the number of topics that matched between a person's own areas of research and what they considered the top three most important areas of research.

***Survey results – scientific articles and journals***

Survey participants were able to write in up to five journals that they felt were the most influential in the field of disease ecology, other than *Science* or *Nature*. Below is a list of the journals that were written in at least twice (note that we only used journals written in four or more times for analyses):

*Proceedings of the Royal Society - Biology (98)*
*Ecology Letters (90)*
*PNAS (85)*
*Ecology (72)*
*Journal of Animal Ecology (60)*
*American Naturalist (44)*
*Trends in Ecology and Evolution (40)*
*Plos Neglected Tropical Diseases (35)*
*Plos Pathogens (33)*
*Journal of Wildlife Diseases (31)*
*Emerging Infectious Diseases (30)*
*Plos ONE (27)*
*International Journal for Parasitology (25)*
*Parasitology (25)*
*Plos Biology (18)*
*EcoHealth (17)*
*Evolution (16)*
*Plos Computational Biology (16)*
*Trends in Parasitology (15)*
*American Society of Tropical Medicine and Hygiene (14)*
*Parasites and Vectors (14)*
*Journal of Parasitology (13)*
*Functional Ecology (12)*
*Journal of Medical Entomology (12)*
*Molecular Ecology (12)*
*Vector-Borne and Zoonotic Diseases (11)*
*Epidemics (9)*
*Philosophical Transactions of the Royal Society B (8)*
*Diseases of Aquatic Organisms (7)*
*eLife (7)*
*Epidemiology and Infection (7)*
*Journal of the Royal Society Interface (7)*
*Journal of Wildlife Management (5)*
*Conservation Biology (5)*
*American Journal of Epidemiology (5)*
*Journal of Infectious Diseases (5)*
*Journal of Applied Ecology (4)*
*Journal of Vector Ecology (4)*
*Journal of Virology (4)*
*Lancet Infectious Diseases (4)*

*Nature: Ecology and Evolution (4)*
*Journal of Evolutionary Biology (4)*
*Ecological Applications (3)*
*Ecological Modeling (3)*
*Frontiers in Ecology and the Environment* (3)
*Journal of Theoretical Biology (3)*
*Malaria Journal (3)*
*Oikos (3)*
*Preventive Veterinary Medicine (3)*
*Biology Letters (2)*
*Biomed Central Biology (2)*
*Cell (2)*
*Ecography (2)*
*Evolutionary Applications (2)*
*Genome Medicine (2)*
*Infection Genetics and Evolution (2)*
*International Society for Microbial Ecology (2)*
*Journal of Clinical Microbiology (2)*
*Landscape Ecology (2)*
*Methods in Ecology and Evolution (2)*
*Oecologia (2)*
*Peer Community in Ecology (2)*
*Peer Community in Evolution (2)*
*Plos Genetics (2)*
*Transboundary and Emerging Diseases (2)*
*Trends in Microbiology (2)*
*Veterinary Parasitology (2)*
*Zoonoses and Public Health (2)*

Respondents could also write in up to five journal articles that were the most influential to their career as a disease ecologist (Table S4).

**Table S4**. Articles that survey respondents considered most influential to their career as a disease ecologist and the number of times each article was written in (n). Note that this table only includes articles written three or more times.

| n | article |
|---|---|
| 18 | Hudson et al. 1998. Prevention of Population Cycles by Parasite Removal. *Science* |
| 13 | Anderson and May. 1979. Population biology of infectious diseases: Part I. *Nature* |
| 12 | Anderson & May. 1978. Regulation and Stability of Host-Parasite Population Interactions: I. Regulatory Processes. *Journal of Animal Ecology* |
| 10 | Lloyd-Smith et al. 2005. Superspreading and the effect of individual variation on disease emergence. *Nature* |
| 9 | Keesing et al. 2006. Effects of species diversity on disease risk. *Ecology Letters* |
| 8 | Jones et al. 2008. Global trends in emerging infectious diseases. *Nature* |
| 8 | Lloyd-Smith et al. 2005. Should we expect population thresholds for wildlife disease? *Trends in Ecology & Evolution* |
| 8 | Lloyd-Smith et al. 2009. Epidemic dynamics at the human-animal interface. *Science* |
| 6 | Daszak et al. 2000. Emerging infectious diseases of wildlife -- threats to biodiversity and human health. *Science* |
| 6 | Grenfell et al. 2004. Unifying the epidemiological and evolutionary dynamics of pathogens. *Science* |
| 6 | Pedersen and Fenton 2007. Emphasizing the ecology in parasite community ecology. *Trends in Ecology & Evolution* |
| 5 | Altizer et al 2003. Social organization and parasite risk in mammals: integrating theory and empirical studies. *Annual Review of Ecology, Evolution, and Systematics* |
| 5 | Grenfell et al. 2001. Travelling waves and spatial hierarchies in measles epidemics. *Nature* |
| 5 | Kuris et al. 2008. Ecosystem energetic implications of parasite and free-living biomass in three estuaries. *Nature* |
| 5 | May & Anderson. 1978. Regulation and Stability of Host-Parasite Population Interactions: II. Destabilizing Processes. *Journal of Animal Ecology* |
| 5 | Plowright et al. 2017. Pathways to zoonotic spillover. *Nature Reviews Microbiology* |
| 4 | Altizer et al. 2006 Seasonality and the dynamics of infectious diseases. *Ecology Letters* |

| | |
|---|---|
| 4 | Altizer et al. 2011. Animal migration and infectious disease risk. *Science* |
| 4 | Altizer et al. 2013. Climate change and infectious diseases: from evidence to a predictive framework. *Science* |
| 4 | Anderson and May. 1981. The population dynamics of microparasites and their invertebrate hosts. *Philosophical Transactions of the Royal Society B* |
| 4 | Dobson. 2004. Population dynamics of pathogens with multiple host species. *The American Naturalist* |
| 4 | Keesing et al. 2010. Impacts of biodiversity on the emergence and transmission of infectious diseases. *Nature* |
| 4 | May and Anderson. 1979. Population biology of infectious diseases II. *Nature* |
| 4 | Sheldon and Verhulst. 1996. Ecological immunology: costly parasite defences and trade-offs in evolutionary ecology. *Trends in Ecology & Evolution* |
| 3 | Anderson and May. 1982. Coevolution of hosts and parasites. *Parasitology* |
| 3 | Bjørnstad et al. 2002. Dynamics of measles epidemics: estimating scaling of transmission rates using a time series SIR model. *Ecological Monographs* |
| 3 | Davies and Pedersen. 2008 Phylogeny and geography predict pathogen community similarity in wild primates and humans. *Proceedings of the Royal Society B* |
| 3 | Fenton and Pedersen. 2005 Community epidemiology framework for classifying disease threats. *Emerging Infectious Diseases* |
| 3 | Graham et al. 2010. Fitness correlates of heritable variation in antibody responsiveness in a wild mammal. *Science* |
| 3 | Haydon et al. 2002. Identifying reservoirs of infection: a conceptual and practical challenge. *Emerging Infectious Diseases* |
| 3 | Hudson et al. 2002. Ecology of wildlife diseases |
| 3 | Hudson et al. 2006. Is a healthy ecosystem one that is rich in parasites? *Trends in Ecology & Evolution* |
| 3 | Levin. 1992. The problem of pattern and scale in ecology. *Ecology* |
| 3 | Lips et al. 2006. Emerging infectious disease and the loss of biodiversity in a Neotropical amphibian community. *Proceedings of the National Academy of Sciences* |
| 3 | Mordecai et al. 2013. Optimal temperature for malaria transmission is dramatically lower than previously predicted. *Ecology Letters* |
| 3 | Råberg et al. 2007. Disentangling genetic variation for resistance and tolerance to infectious diseases in animals. *Science* |
| 3 | Råberg et al. 2008. Decomposing health: tolerance and resistance to parasites in animals. *Philosophical Transactions of the Royal Society B* |
| 3 | Restif et al. 2012. Model-guided fieldwork: practical guidelines for multidisciplinary research on wildlife ecological and epidemiological dynamics. *Ecology Letters* |
| 3 | Telfer et al. 2010. Species interactions in a parasite community drive infection risk in a wildlife population. *Science* |

**Literature Search Methods**

The literature search was conducted in Web of Science for the years 1975 to 2018. Each article had to meet specific criteria using Boolean filters: (1) an indicator that a pathogen or parasite is being studied (virus* OR *pathogen* OR bacteria OR *parasite* OR infectio*); (2) an indicator that the focus of the article was among individuals (e.g. not cellular level) (population* OR *system* OR transmission); and (3) an indicator that the host species falls within the remit of disease ecology (*host* OR human* OR animal* OR wildlife OR *vector*). We used exclusionary terms to remove groups of similar but non-disease ecology articles; specifically, we excluded 'brood parasite' OR 'parasitic wasp' OR sequence OR uptake OR polymorphism* OR 'side effects' OR protein* OR receptor OR isolate* or HeLa OR maker* OR *methyl*. We used

Web of Science categories to narrow our search after deducing that they reduced our false-positive rate. Web of Science categories included ecology, entomology, infectious diseases, mathematical and computational biology, biology, evolutionary biology, fisheries, plant sciences, marine and freshwater biology, tropical medicine, genetics and heredity, parasitology, multidisciplinary sciences, veterinary sciences, virology, and public, environmental, occupational, or health. Below is the full search in Web of Science. Document types were then narrowed to articles, reviews, Proceedings papers, book reviews, editorial material (e.g., short articles), notes, and letters (e.g., comments about an article). Finally, articles with fewer than four citations were removed as a form of quality control.

TOPICS:
(virus* OR *pathogen* OR bacteria OR *parasite* OR infectio*) AND
(population* OR *system* OR transmission) AND
(*host* OR human* OR animal* OR wildlife OR *vector*)
NOT:
('brood parasite' OR 'parasitic wasp' OR sequence OR uptake OR polymorphism* OR 'side effects' OR protein* OR receptor OR isolate* or HeLa OR maker* OR *methyl*)

JOURNALS (44):
(SCIENCE) OR
(NATURE) OR
(PROCEEDINGS OF THE ROYAL SOCIETY B-BIOLOGICAL SCIENCES) OR
(ECOLOGY LETTERS) OR
(PROCEEDINGS OF THE NATIONAL ACADEMY OF SCIENCES OF THE UNITED STATES OF AMERICA) OR
(ECOLOGY) OR
(JOURNAL OF ANIMAL ECOLOGY) OR
(AMERICAN NATURALIST) OR
(TRENDS IN ECOLOGY & EVOLUTION) OR
(PLOS NEGLECTED TROPICAL DISEASES) OR
(PLOS PATHOGENS) OR
(JOURNAL OF WILDLIFE DISEASES) OR
(EMERGING INFECTIOUS DISEASES) OR
(PLOS ONE) OR
(INTERNATIONAL JOURNAL FOR PARASITOLOGY) OR
(PARASITOLOGY) OR
(PLOS BIOLOGY) OR
(ECOHEALTH) OR
(EVOLUTION) OR
(PLOS COMPUTATIONAL BIOLOGY) OR
(TRENDS IN PARASITOLOGY) OR
(AMERICAN JOURNAL OF TROPICAL MEDICINE "AND" HYGIENE) OR
(PARASITES "AND" VECTORS) OR
(JOURNAL OF PARASITOLOGY) OR
(FUNCTIONAL ECOLOGY) OR
(JOURNAL OF MEDICAL ENTOMOLOGY) OR
(MOLECULAR ECOLOGY) OR
(VECTOR-BORNE "AND" ZOONOTIC DISEASES) OR

(EPIDEMICS) OR
(PHILOSOPHICAL TRANSACTIONS OF THE ROYAL SOCIETY B-BIOLOGICAL SCIENCES) OR
(DISEASES OF AQUATIC ORGANISMS) OR
(ELIFE) OR
(EPIDEMIOLOGY "AND" INFECTION) OR
(JOURNAL OF THE ROYAL SOCIETY INTERFACE) OR
(JOURNAL OF WILDLIFE MANAGEMENT) OR
(CONSERVATION BIOLOGY) OR
(AMERICAN JOURNAL OF EPIDEMIOLOGY) OR
(JOURNAL OF INFECTIOUS DISEASES) OR
(JOURNAL OF APPLIED ECOLOGY) OR
(JOURNAL OF VECTOR ECOLOGY) OR
(JOURNAL OF VIROLOGY) OR
(LANCET INFECTIOUS DISEASES) OR
(ECOLOGY "AND" EVOLUTION) OR
(JOURNAL OF EVOLUTIONARY BIOLOGY)

WEB OF SCIENCE CATEGORIES:
ECOLOGY OR
ENTOMOLOGY OR
INFECTIOUS DISEASES OR
MATHEMATICAL & COMPUTATIONAL BIOLOGY OR
BIOLOGY OR
EVOLUTIONARY BIOLOGY OR
FISHERIES OR
PUBLIC ENVIRONMENTAL OCCUPATIONAL HEALTH
OR PLANT SCIENCES OR
MARINE & FRESHWATER BIOLOGY OR
TROPICAL MEDICINE OR
GENETICS & HEREDITY OR
PARASITOLOGY OR
MULTIDISCIPLINARY SCIENCES OR
VETERINARY SCIENCES OR
VIROLOGY

YEARS 1975-2018

ALL PUBLICATIONS WITH ABSTRACTS (AVAILABLE in ENGLISH)

WoS Core Collection: Science Citation Expanded 1900—Present

DOWNLOADED FEBRUARY 4, 2020

**False-Positive Literature Assessment**

To identify and quantify the false-positive rate of our literature corpus, two authors (DJB and KMF) independently evaluated the same 100 randomly selected articles and classified them as 'disease ecology' or 'outside the field'. Both authors classified 75/100 papers as disease ecology; the 75 'disease ecology' and 25 'outside the field' papers were very similar (94% agreement, Cohen's κ=0.84), with both authors classifying the same 22 papers as 'outside the field'. False-positive studies were primarily about pathogenesis or microbiology. We did not detect an association between topical and temporal trends ($\chi^2$ = 72.84, p = 0.29). Figure S5 shows the type of study and year published of the false-positive papers; note that although the number of false-positives increases in more recent years, the total number of publications substantially increased over this period (i.e., since 2000).

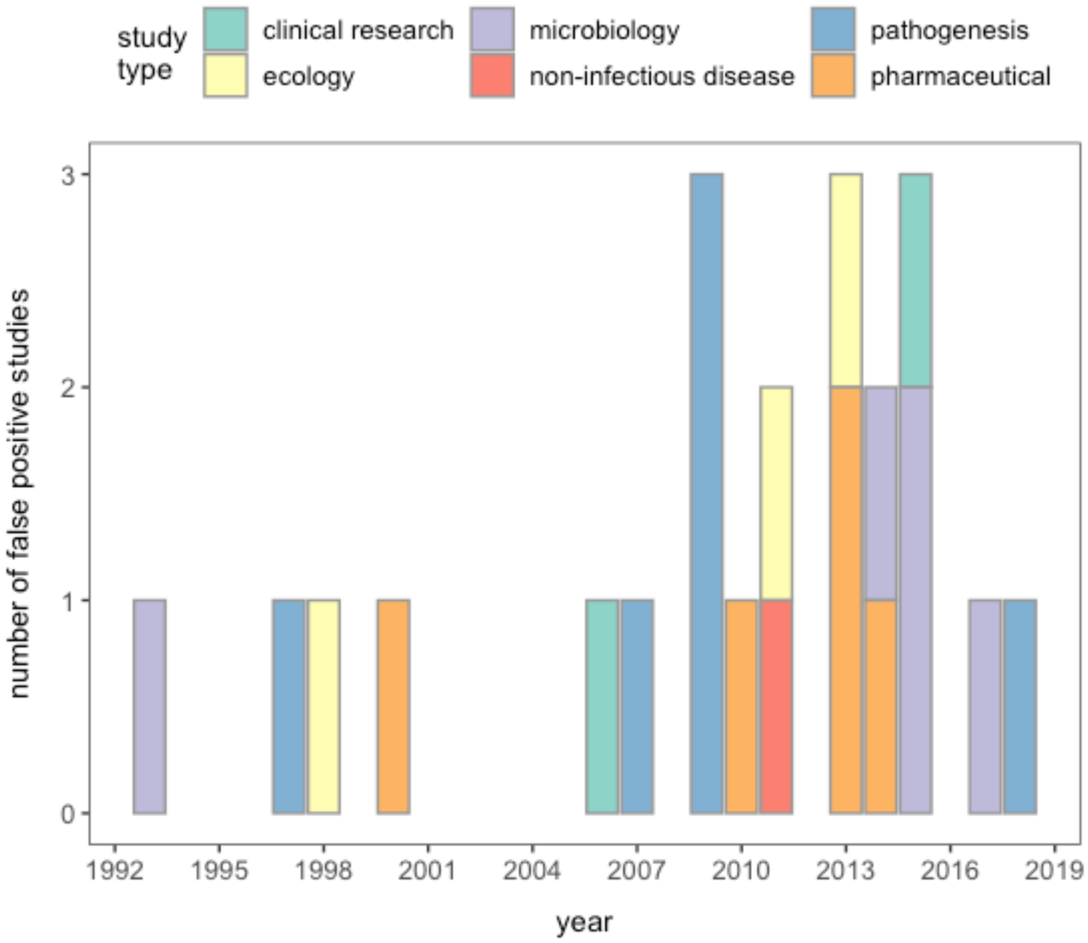


**Figure S5**. A count of the papers classified as "outside the field" of disease ecology based on their year of publication (n=22). Papers are colored by study type.

**Additional Literature Review Results**

Below are summary outputs from the Web of Science database application that provide a more comprehensive overview of the literature corpus.

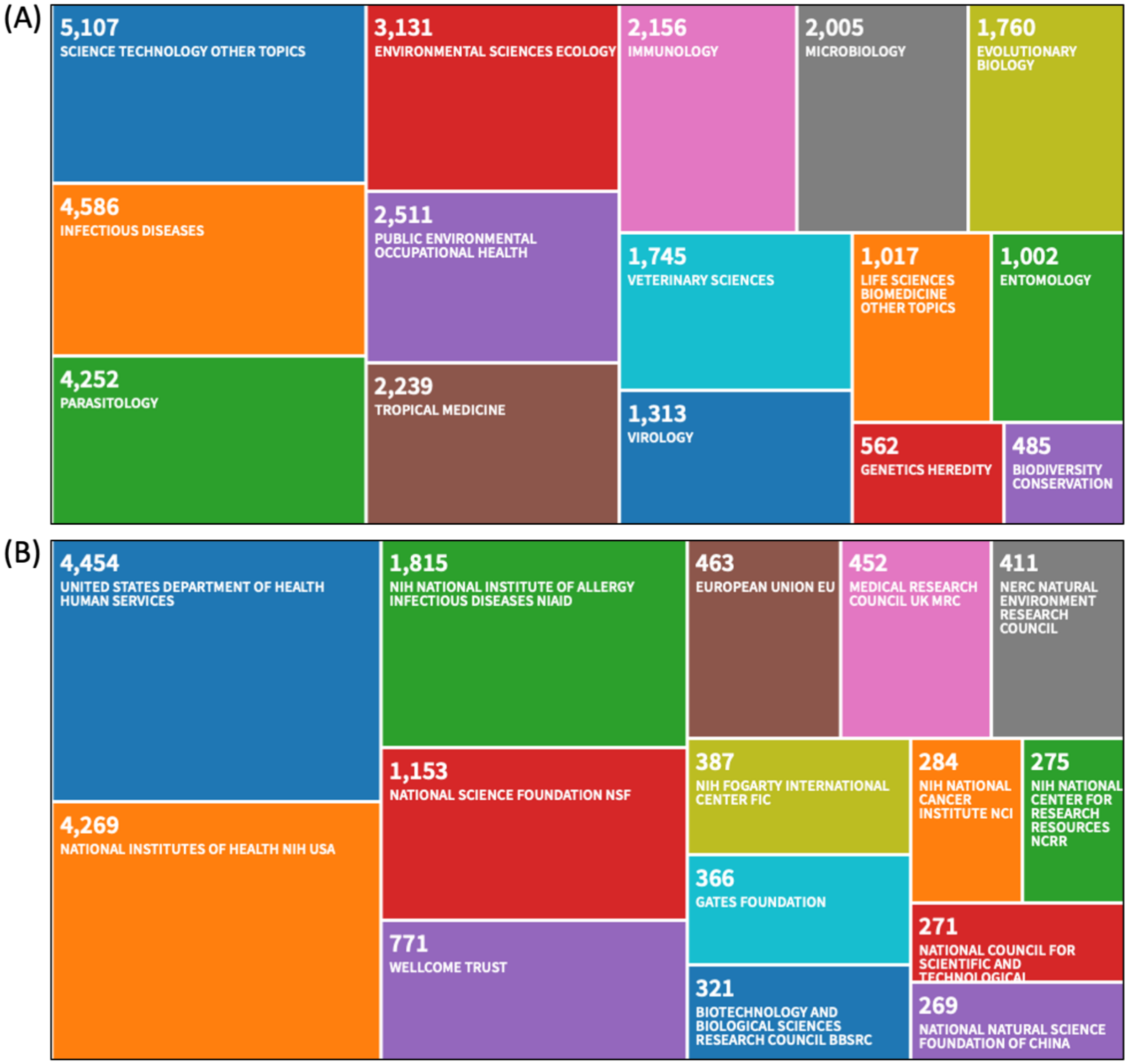


**Figure S6**. Treemap of the top 15 (A) research areas and (B) funding institutions reported in the literature corpus (Web of Science, April 2020).

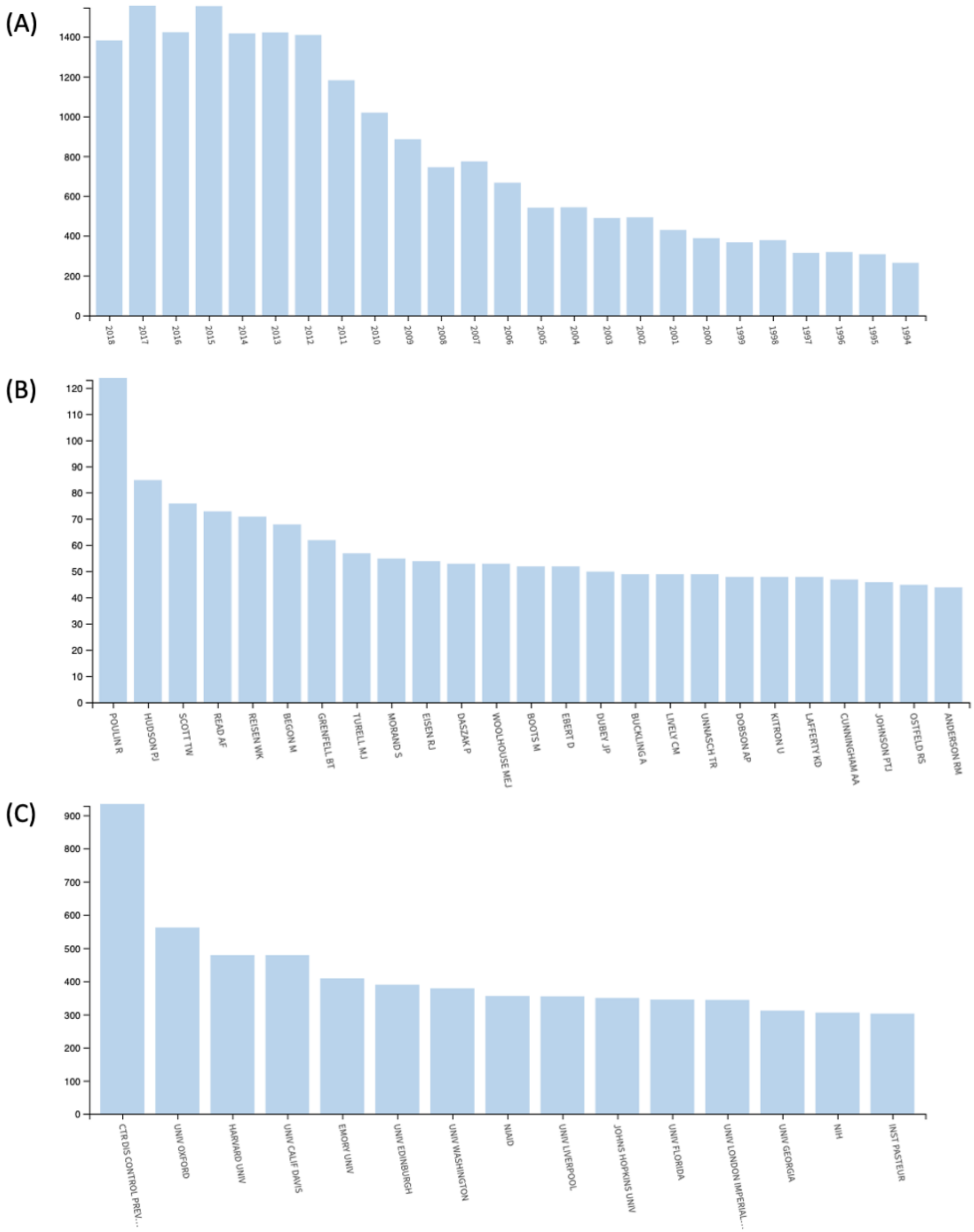


**Figure S7**. Histograms displaying the count of (A) published papers over the last 25 years, from the most recent year in the corpus (2018) to 1994; (B) publications in the corpus for the top 25 authors (names are written as: LAST “space” FIRST [AND MIDDLE] INITIAL); (C) publications in the corpus by the top 15 organizations (Web of Science, April 2020).

***Topic detection – emerging from the literature***

Below are the 15 topic clusters over the entire corpus and assigned names. Confidence in naming each topic was on a scale from 1 (high confidence) to 3 (low confidence). Overall, we had high confidence in ascribing a name to most topics. Low confidence arose in topics with many general words, especially wildlife pathogens (Topic 12) and infectious disease modeling (Topic 1). To assess trends statistically and visually (e.g., Fig. 4B), words with very little predictive value were removed (e.g., infection, virus, data, host, human, etc.) because these words were too common to help distinguish unique trends.

Topic #1: INFECTIOUS DISEASE MODELING (confidence = 2)
transmission, risk, model, human, case, outbreak, health, control, epidemic, data, infectious, animal, factor, contact, area

Topic #2: INFECTION TRIALS, GENETICS (confidence = 1)
cell, human, viral, expression, gene, replication, tissue, type, vitro, vivo, infection, infected, culture, macrophage, blood,

Topic #3: EVOLUTION OF VIRULENCE (fish hosts) (confidence = 2)
parasite, host, parasitology, fish, transmission, intermediate, population, infected, stage, evolution, life, interaction, virulence, parasitic, nematode

Topic #4: MOSQUITO-BORNE PATHOGENS (confidence = 1)
mosquito, vector, aedes, wnv, ae, aegypti, dengue, culex, nile, west, transmission, wolbachia, female, feeding, temperature

Topic #5: HIV (confidence = 1)
hiv, immunodeficiency, woman, patient, men, therapy, drug, human, infectious, transmission, aids, treatment, background, risk, type

Topic #6: SEROLOGY (confidence = 1)
infection, prevalence, antibody, sample, infected, age, population, human, patient, positive, animal, serum, detected, associated, child

Topic #7: IMMUNE RESPONSE (confidence = 1)
response, immune, immunity, system, infection, cytokine, antigen, activation, systemic, mechanism, function, level, model, induced, expression

Topic #8: TROPICAL MOSQUITO-BORNE PATHOGENS (confidence = 1)
background, neglected, tropical, dengue, fever, human, aedes, control, africa, burden, factor, country, aegypti, vector, treatment

Topic #9: TICK-BORNE PATHOGENS IN RODENT HOSTS (confidence = 2)
tick, vector, deer, adult, feeding, larva, pathogen, host, transmission, infected, collected, site, medical, fever, rodent

Topic #10: MALARIA (confidence = 1)
malaria, falciparum, plasmodium, mosquito, child, parasite, blood, culture, africa, human, vector, drug, development, stage, control

Topic #11: INFLUENZA (confidence = 1)
influenza, avian, bird, respiratory, pathogenic, seasonal, strain, human, viral, highly, transmission, epidemic, wild, infectious, pig

Topic #12: “GENERIC” WILDLIFE PATHOGENS (confidence = 3)
species, community, host, diversity, abundance, bird, amphibian, distribution, prevalence, habitat, fish, site, wild, pattern, population

Topic #13: INFECTION TRIALS [in mice and other species] (confidence = 2)
mouse, model, strain, infection, brain, day, infected, systemic, vivo, animal, treatment, human, pathogenesis, deer, gene

Topic #14: VACCINE RESEARCH AND DEVELOPMENT (confidence = 1)
vaccine, vaccination, hpv, protection, efficacy, trial, challenge, immunity, human, strain, cancer, development, effective, antibody, type

Topic #15: PLANT/HOST-PATHOGEN COEVOLUTION (confidence = 1)
pathogen, host, resistance, plant, population, evolution, gene, interaction, genetic, dynamic, virulence, strain, bacterial, bacteria, model

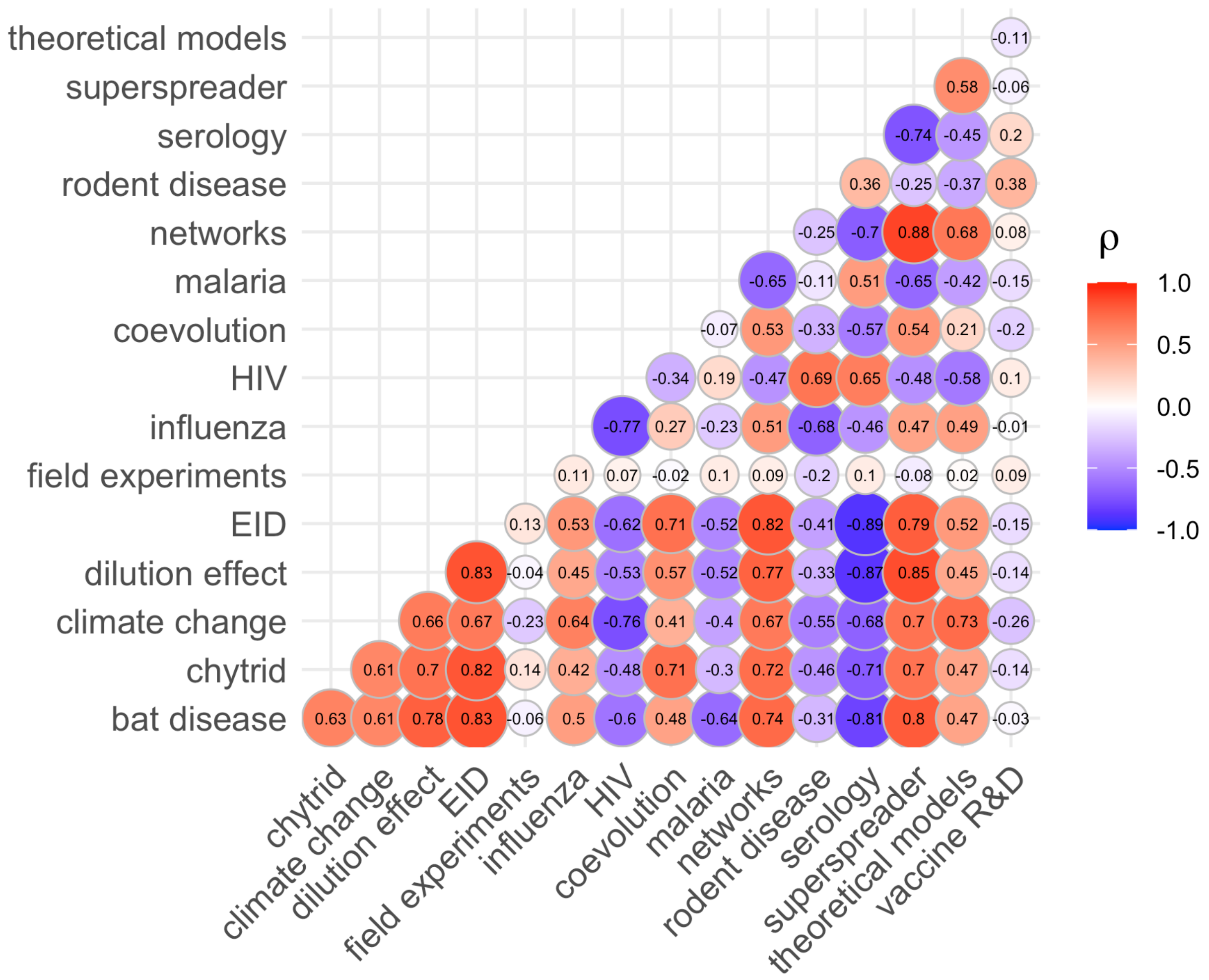


**Figure S8**. Spearman's correlation coefficients (ρ) between the proportion of publications per topic over time. Red indicates positive covariation between two time series, whereas purple indicates negative covariation. Circles are sized according to the absolute value of the correlation.

***Topic detection – selected by authors***

Below are the author selected topics we assessed.

Emerging infectious diseases (EID):
emerge, zoonosis, spillover, outbreak, reservoir, epidemic, pandemic, contain

Climate change:
climate, change, range, distribution, biogeography, landscape, global, transmission, temperature

Network models:
network, contact, social, edge, node, centrality, connectivity, epidemiology, dynamic, transmission

Dilution Effect:

dilution, amplification, hypothesis, diversity, community, risk, transmission, infection, effect

Neutral:
analysis, study, paper

Rodent:
rodent, mouse, rat

Bat:
bat, Chiroptera

Chytrid:
chytrid, Chytridiomycosis, amphibian, fungus

Superspreader:
superspreader, epidemic, contact, variation

Theoretical models:
SIR, theoretical, model, differential equation, R0, mathematical

Field experiments:
field, experiment, treatment, factorial, control
